# VGF in the Nucleus Accumbens Regulates Synaptic and Opioid-Evoked Plasticity

**DOI:** 10.64898/2025.12.10.693464

**Authors:** Anisha P. Adke, Lydia B. Cunitz, Rachel M. Dick, Tuba H. Ali, Carolina A. M. Rocha, Ying Zhang, Lucy Vulchanova, Patrick E. Rothwell

## Abstract

Neuropeptides contribute to the functional complexity of the nucleus accumbens, and evaluating the mechanisms by which they influence reward neurocircuitry may reveal novel therapeutic targets. In this study, we investigated the role of the neuropeptide precursor VGF (non-acronymic) in the nucleus accumbens, and its contribution to synaptic and opioid-evoked plasticity and behavior. We first characterized VGF expression within the nucleus accumbens, then examined its impact on excitatory synaptic transmission, and finally disrupted its expression to assess its influence on opioid sensitivity. We found that VGF is expressed by interneurons and medium spiny neurons in the nucleus accumbens, and the VGF-derived peptide TLQP-62 reduced excitatory synaptic transmission onto nucleus accumbens medium spiny neurons. Conditional genetic knockout of VGF from the nucleus accumbens did not affect acute opioid sensitivity, but amplified opioid-evoked psychomotor sensitization following chronic fentanyl exposure. These results establish VGF as a modulator of nucleus accumbens synaptic and drug-evoked plasticity. Uncovering the impact of VGF on nucleus accumbens plasticity following exogenous opioid exposure may lead to novel therapeutic targets to treat opioid use disorders.

**SIGNIFICANCE STATEMENT:** The opioid crisis has continued to have a devastating impact, and understanding the basic mechanisms underlying opioid use disorder may facilitate the discovery of new therapeutic targets. Neuropeptide signaling in the nucleus accumbens may be a potential site for intervention given the state-dependent recruitment of neuropeptides in reward circuitry. The neuropeptide precursor VGF has never been studied within the context of drug-evoked plasticity and behavior, despite its role in both adaptive and maladaptive plasticity, and evidence that drug exposure modulates its expression. Our findings show that VGF may be a novel modulator of NAc synaptic and opioid-evoked plasticity, building our understanding of the mechanisms responsible for drug-evoked adaptations and identifying VGF as an interesting potential target for future investigation.

## INTRODUCTION

The nucleus accumbens (NAc) is a brain region critical for processing reward, motivation, and affect, integrating complex information and translating it into behavioral action. Within the NAc, neuropeptides tune excitability and synaptic dynamics to adjust behavioral responses to changing environmental and internal states (Castro and Bruchas, 2019). Peptide signaling can shape how NAc neurons encode reward (Castro and Berridge, 2014; Al-Hasani et al., 2015; Castro et al., 2016), salience (Warthen et al., 2019), and affective information (Lemos et al., 2012; Thorsell and Mathé, 2017), thereby guiding adaptive behavior. In the context of opioid use disorders, exposure to exogenous opioids can topple the intricate balance of neuropeptide signaling, disrupting plasticity and altering motivational states (Nylander et al., 1995; Trujillo et al., 1995; Solecki et al., 2009; Schlosburg et al., 2013; Atwood et al., 2014). In turn, neuropeptides influence the physiological and behavioral effects of drugs, modulating the circuitry that governs reward-related behaviors (Lu et al., 2003; Chung et al., 2009; Cohen et al., 2014; Ehrich et al., 2014; Nam et al., 2019). Neuropeptide signaling is therefore a critical regulator of NAc physiology and maladaptive responses to addictive substances.

While some peptides have been extensively studied for their alterations in pathological states, the contribution of the neuropeptide precursor VGF (non-acronymic) to NAc physiology and reward-related behavior is not well studied. VGF is broadly expressed across the nervous system and is cleaved into numerous bioactive peptides, most of which have undefined roles (Trani et al., 2002; Levi et al., 2004; Ferri et al., 2011; Quinn et al., 2021). Among these, the C-terminal peptide TLQP-62 (named after the first 4 amino acids and length of the sequence) has been linked to the regulation of synaptic plasticity and excitatory neurotransmission in the hippocampus and spinal cord (Alder et al., 2003; Bozdagi et al., 2008; Moss et al., 2008; Skorput et al., 2018). VGF expression levels in the NAc are altered across numerous physiological and pathological contexts, including stress and mood-related states (Jiang et al., 2018) as well as exposure to addictive substances (Heiman et al., 2008; Noli et al., 2017; Wimmer et al., 2019; Lefevre et al., 2020), suggesting that VGF may serve as a flexible neuromodulatory signal within mesolimbic circuitry. However, without a clear understanding of which NAc cell populations express VGF, how VGF-derived peptides influence NAc physiology, and which behavioral processes rely on VGF signaling, the mechanistic basis for these observations remains unknown.

Here, we take a multi-level approach to characterize VGF within the NAc and to define its functional relevance to NAc circuitry and opioid-evoked behaviors. We first performed a cross-species meta-analysis of gene expression data from mouse, rat, and humans to identify the NAc cell populations that express VGF. We then validated and expanded upon these data by mapping the spatial organization of VGF expression in the NAc with fluorescent in situ hybridization. Using whole-cell patch-clamp recordings, we next measured the acute physiological effects of the VGF-derived peptide TLQP-62 on synaptic transmission in NAc medium spiny neurons (MSNs). Finally, we used a conditional genetic knockout to selectively disrupt VGF expression, and examined the impact on behavioral responses to acute and chronic opioid administration. Together, these experiments define cellular, synaptic, and behavioral roles of VGF in the NAc and establish this peptide precursor as an important and understudied modulator of reward-related neurocircuitry.

## METHODS

### RNA sequencing meta-analysis

Publicly available single-cell and single-nucleus RNA-sequencing datasets derived from mouse, rat, and human nucleus accumbens were retrieved from previously published studies (Avey et al., 2018; Saunders et al., 2018; Savell et al., 2020; Chen et al., 2021; Tran et al., 2021). Raw counts or processed gene-expression matrices were downloaded from authors’ repositories and processed in R (v4.4.2) and Seurat (v5.3.0) following standard workflows. In studies assessing the effects of a drug manipulation, only control datasets were used. For each dataset, we replicated the preprocessing and clustering pipelines described in the original publications to reproduce published cluster assignments. Because the cell-type annotations varied across studies in terms of resolution, we collapsed clusters into a common set of broad, conserved cell-type categories (e.g., MSNs, inhibitory interneurons, astrocytes, oligodendrocytes, microglia) to enable valid cross-study comparisons. Normalized log transformed expression values for VGF were then extracted and merged across datasets, and dot-plot visualizations were generated to depict cell-type-specific expression patterns.

### Animals

All procedures were approved by the Institutional Animal Care and Use Committee at the University of Minnesota and followed the NIH Guidelines for the Care and Use of Laboratory Animals. Male and female C57BL/6J mice (Strain #000664) were obtained from The Jackson Laboratory. The conditional VGF knockout mouse line (*Vgf^flpflox/flpflox^*, abbreviated VGF^fl/fl^ throughout the remainder of the text; Strain #030571, The Jackson Laboratory) was a gift from Stephen Salton (Jiang et al., 2018) and was maintained on a C57Bl/6J background. Mice used for electrophysiology carried one copy of the Drd1a-tdTomato bacterial artificial chromosome (BAC) transgene (Shuen et al., 2008) and/or the Drd2-eGFP BAC transgene (Gong et al., 2003). Mice were 5-8 weeks old at the beginning of each experiment and housed in groups of 2-5. Cages were kept at ∼23°C on a 12-hour light cycle (0600h - 1800h). Food and water were provided *ad libitum*. All experiments were performed between 800h and 1700h, with behavior studies occurring in the morning between 800h and 1300h.

### RNA fluorescent in situ hybridization (FISH)

Mice were deeply anesthetized using isofluorane, and depth of anesthesia was confirmed with a hard toe pinch. Perfusions were performed through the heart using cold calcium-free Tyrode’s solution (in mM: 116 NaCl, 5.4 KCl, 1.6 MgCl_2_●6H_2_O, 0.4 MgSO4●7H2O, 1.4 NaH_2_PO_4_, 5.6 glucose, and 26 Na_2_HCO_3_) until the liver cleared, followed by ∼200 mL of cold fixative (4% paraformaldehyde, pH 6.9). Brains were extracted and post-fixed overnight at 4°C, then placed in 10% sucrose for at least 24 hours. One hour prior to cryosectioning, brains were rapidly frozen using pressurized CO_2_ for approximately 1 minute in OCT. Following equilibration in the cryostat (−18°C), 14 μm sections were cut and placed onto SuperFrost Plus microscope slides (Fisher Brand) and stored at either −20°C (for less than 4 months) or −80°C (for greater than 4 months). RNA FISH was performed using the Advanced Cell Diagnostics RNAscope Multiplex Fluorescent V2 Assay, with methods modified from the user manual. Unless specified, all reagents were from ACD Bio.

In brief, slides were postfixed in cold 4% paraformaldehyde, pH 6.9 for 10 minutes then dehydrated progressively using 50% (5 minutes), 70% (5 minutes), and 100% (2 x 5 minutes) ethanol. After removing from 100% ethanol and drying, several drops of hydrogen peroxide were added to the slide to fully cover brain sections, followed by incubation for 10 minutes at room temperature. Slides were rinsed in double distilled water (ddH_2_O) three times and then incubated in Target Retrieval Reagent for 4 minutes at 92-97°C. Slides were immediately transferred to room temperature ddH2O, rinsed twice, then placed in 100% ethanol for 5 minutes. After slides dried, a hydrophobic barrier was drawn around the sections and dried for ∼10 minutes at room temperature. Protease III was added to the sections, and slides were incubated for 15 minutes at 40°C in a humidified box placed in an incubator. Slides were rinsed twice for 2 minutes in ddH_2_O, then excess fluid was flicked off, and probes previously warmed to 40°C and equilibrated to room temperature were added. The probes used were Mm-Vgf-C1 (Cat No. 517421), Mm-Sst-C2 (Cat No. 404631-C2), Mm-Drd1-C2 (Cat. No. 461901-C2), and Mm-Drd2-C3 (Cat. No. 406501-C3). Slides were incubated with the probes for 2 hours at 40°C in a humidified box, then washed twice on a shaker for 2 minutes in 1x RNAscope Wash Buffer. Amplification steps were then performed at 40°C in a humidified box using AMP1 for 30 minutes, AMP2 for 30 minutes, and AMP3 for 15 minutes, with two washes performed between each amplification step.

For VGF and Sst co-labeling studies, AMP4 Alt B was added for 45 minutes at 40°C, rinsed twice in Wash Buffer, labeled with DAPI for 10 seconds, and coverslipped using Fluorosave (Millipore Sigma). For VGF, Drd1, and Drd2 co-labeling studies, each channel was opened and labeled in the following order, all at 40°C: HRP-1 for 15 minutes; Opal-570 (Akoya Biosciences, 1:3000) diluted in Tyramide Signal Amplification (TSA) for 20 minutes; HRP Blocker for 15 minutes; HRP-2 for 15 minutes; Opal-520 (Akoya Biosciences, 1:3000 diluted in TSA) for 15 minutes; HRP Blocker for 15 minutes; HRP-3 for 15 minutes; Opal-690 (Akoya Biosciences, 1:2000 diluted in TSA) for 15 minutes; and HRP Blocker for 15 minutes. Slides were washed twice for 2 minutes in Wash Buffer between each step. After applying DAPI for 10 seconds, slides were rinsed briefly in wash buffer and coverslipped with Fluorosave (Millipore Sigma).

### Confocal imaging and FISH analysis

Images were captured digitally using a Nikon A1RSI Confocal with SIM Super Resolution (objective: 20x, NA 0.75) and Nikon NIS Elements software (v. 5.30.05, Nikon, Melville, NY). Nyquist zoom (9.61, 0.13 pixel sampling rate) was used, and 5 x 5 images were stitched to sample a larger field of view. At least six regions in the NAc (dorsal, ventral, medial, and lateral; core and shell) were sampled within one hemisphere. Laser intensity was manually adjusted for each image, with the DAPI channel being overexposed to aid in nuclei identification during analysis. At least 3 slices per animal (6 animals for VGF/Drd1/Drd2 analysis, 8 animals for VGF/Sst analysis) were used. For anatomical analysis of VGF expression, the core, shell, and anterior-posterior (AP) location were identified using a brain atlas (Paxinos and Franklin, 2001). Anterior NAc was defined as AP +1.54-1.98, middle as +1.1-1.42, and posterior as +0.62-0.98.

Analysis was performed using HALO Pathology Image Analysis Software (v3.6.4134, Indica Labs). For VGF/Sst analysis, we used the algorithm HighPlex FL (v4.2, Indica Labs) to identify nuclei and detect transcripts. Because different fluorophores were used for VGF/Drd1/Drd2, we used the FISH algorithm (v.3.0.4, Indica Labs). DAPI expression was used to detect nuclei. Contrast threshold and minimum signal intensity were adjusted for each probe based on brightness and background signal. FISH scoring was used to calculate degrees of expression (low, medium, high) using percent of coverage (for VGF/Sst) and number of transcript copies (for VGF/Drd1/Drd2). On average, cells containing more than 4 individual transcript copies were counted as positive for VGF, Sst, Drd1, or Drd2. VGF expression was low if the cell had 5-10 transcript copies, medium if 10-15 transcript copies, and high if it contained over 15 transcript copies. Settings were manually adjusted for each image and any tissue imperfections were excluded from the data.

### Western blots

Mice were deeply anesthetized using isoflurane. Brains were extracted and coronally sectioned using a brain matrix (1 mm thick). Sections containing the NAc were microdissected using a 1 mm circular tissue biopsy punch (Electron Microscopy Sciences), resulting in 4 tissue punches containing bilateral accumbens for each animal. The tissue was immediately flash-frozen using dry ice and stored at −20°C until used. Western blots were performed as previously described (Skorput et al., 2018). Tissue was homogenized in lysis buffer (Tris-buffered saline, pH 7.4, containing 1% Triton X-100, 10 mM EGTA, and 10 mM EDTA) with protease inhibitors (EDTA-free Complete Mini, Roche) using a glass bead homogenizer (0.5 mm, Scientific Industries) for 5 minutes. The homogenate was centrifuged for 15 minutes at 14,000 G and 4°C. Supernatant (∼40 μL) was collected, and the concentration was determined using a BCA Protein Assay. A volume that yielded 20 μg of protein was then mixed with sample buffer (NuPAGE 4x sample buffer, Invitrogen), heated for 10 minutes at 95°C, and loaded into a 4-12% gradient gel (NuPAGE 4-12% Bis-Tris Gel, Invitrogen) suspended in MES Running Buffer (NuPAGE MES SDS 20x, Invitrogen). Two lanes were loaded with 3 μL of standards (SeeBlue Plus2 Prestained Standard, Invitrogen). The gel was run at 150 volts for 90-120 minutes, fixed in 10% glutaraldehyde for 1 hour, rinsed with TBS, and then transferred to PVDF membranes (Immobilin-FL, Millipore) using 120 mA at 4°C overnight. PVDF membranes were then placed into blocking buffer (LiCor Odyssey Blocking buffer, Invitrogen) on a shaker for 1 hour at room temperature. Membranes were then incubated in primary antisera (guinea pig anti-AQEE-30 serum; 1:5000) (Riedl et al., 2009) for 48 hours at 4°C on a shaker, rinsed with 0.1% Tween-20 in TBS for 1 hour, then incubated in secondary antisera (1:2000, donkey anti-guinea pig HRP-IgG, #706-035-148, RRID AB_2340447, JacksonImmuno) for 2 hours at 4°C on a shaker. After rinsing with 0.1% Tween-20 in TBS for 1 hour, protein bands were visualized using horseradish peroxidase substrate (Pierce ECL Western Blotting Substrate, Thermo Scientific). Membranes were then imaged using an Amersham Imager 600UV with exposures manually adjusted to avoid saturation. Membranes were then washed in TBS-T, stripped (NuPAGE Stripping Buffer, Invitrogen), and rinsed in TBS-T. The reference antibody (1:5000, rabbit anti-beta-actin-HRP, #5125, Cell Signaling Technology) was added overnight at 4°C on a shaker. Membranes were rinsed with TBS-T and bands visualized as described above.

### Whole-cell patch-clamp electrophysiology

Acute brain slices (240 μm thick) containing the NAc were prepared as previously described (Toddes et al., 2021; Retzlaff and Rothwell, 2022; Trieu et al., 2022; Lefevre et al., 2023). Mice carrying either the Drd1a-tdTomato and/or the Drd2-eGFP transgene were anesthetized with isofluorane and decapitated. The brain was rapidly dissected and placed into oxygenated (95% O_2_/5% CO_2_) icy slurry cutting solution containing (in mM): 228 sucrose, 26 NaHCO_3_, 11 glucose, 2.5 KCl, 1 NaH_2_PO_4_-H_2_O, 7 MgSO_4_-7 H_2_0, and 0.5 CaCl_2_-2 H_2_O, 325-335 mOsm. The olfactory bulbs and cerebellum were removed, the hemispheres cut, and the brain was sectioned coronally using a vibratome (Leica VT 1000S). After sectioning, brain slices recovered for 10 minutes in warm (33°C) artificial cerebral spinal fluid (ACSF, 295-300 mOsm) containing (in mM): 119 NaCl, 26.2 NaHCO_3_, 2.5 KCl, 1 NaH_2_PO_4_-H_2_O, 11 glucose, 1.3 MgSO_4_-7 H_2_O, and 2.5 CaCl_2_-2 H_2_O. Slices equilibrated to room temperature for at least 1 hour before use. For the recordings, slices were transferred to a recording chamber and continuously perfused with room temperature ACSF at 1.5-2 mL/min. All solutions were continuously oxygenated (95% O_2_/5% CO_2_).

Whole-cell voltage-clamp recordings were obtained under visual control using differential interference contrast optics and fluorescence on an Olympus BX51WI microscope. Medium spiny neurons expressing Drd1 (D1-MSNs) or Drd2 (D2-MSNs) were identified by the presence of tdTomato or GFP fluorescence, respectively. Recording electrodes were made from borosilicate glass electrodes (3-5 MΩ) filled with an internal solution (pH 7.3, adjusted with CsOH, 275-285 mOsm) composed of (in mM): 120 CsMeSO_4_, 15 CsCl, 10 TEA-Cl, 8 NaCl, 10 HEPES, 1 EGTA, 5 QX-314, 4 ATP-Mg, and 0.3 GTP-Na. The extracellular ACSF bath contained the GABA_A_ receptor antagonist picrotoxin (50 µM, Tocris) and the sodium channel blocker tetrodotoxin (500 nM, Cayman Chemical) to block the generation of action potentials and isolate miniature excitatory postsynaptic currents (mEPSCs). TLQP-62 (TLQPPASSRRRHFHHALPPARHHPDLEAQARRAQEEADAEERRLQEQEELENYIEHVLLHRP, Cat. #317603, NovoPro Labs) and the scrambled control peptide (SQEDPEELNLHREALAHTAREARHQLHHAPQPERFHRPPVYRESEERQARILRADLAPLHQE, NovoPro Labs) were dissolved in ddH_2_O and, on the recording day, were diluted to 50 nM in ACSF. This concentration of TLQP-62 was selected because it has previously been shown to produce changes in excitatory transmission in acute slice preparations (Skorput et al., 2018). Following a >10-minute baseline in ACSF with picrotoxin and tetrodotoxin, TLQP-62 or the scrambled control peptide was washed on. We quantified mEPSCs 10 minutes following bath application of the peptide, to allow for adequate tissue penetration and activation of possible signaling cascades (Lin et al., 2015; Skorput et al., 2018). Recordings continued for at least 20 minutes after the peptide was added to the slice chamber.

Recordings were performed using a MultiClamp 700B amplifier (Molecular Devices) and Dendrite Digitizer (Sutter Instruments) and acquired using SutterPatch (v2.3) in Igor Pro (8.04, WaveMetrics). Data was filtered at 2 kHz and digitized at 10 kHz. Series resistance was continuously monitored, and experiments were excluded if the resistance exceeded 20 MΩ or changed by more than 20%. Easy Electrophysiology software (v2.8.0, RRID SCR_021190) was used for analysis. Recordings were filtered at 1.5 kHz. An amplitude threshold of 5 pA and a detection criterion of 4.00 were used to identify currents. For each cell, 200 events were analyzed for baseline and peptide conditions.

### Surgical Procedures

Adeno-associated viral vectors (pAAV2retro-hSyn-Cre-P2A-tdTomato and AAV2retro-hSyn-Flp-P2A-tdTomato) (Toddes et al., 2024) were packaged by the University of Minnesota Viral Innovation Core and stored at −80°C until use. VGF^fl/fl^ mice were anesthetized using isofluorane/oxygen (4%/96% for induction; 1.5-2.5%/97.5-98.5% for maintenance). Toe pinches were performed every 5 minutes throughout surgery to ensure anesthesia depth and comfort, and mice were kept on a heating pad to maintain body temperature. Carprofen (10 mg/kg) was subcutaneously injected ∼5 minutes before the incision for pain relief. A small hole was drilled bilaterally above the NAc (AP +1.50, ML ±0.90, DV −4.60). A glass needle was lowered slowly to DV −4.65, then retracted to −4.60. A volume of 500 nL of virus was delivered to each side at a rate of 0.1 μL/min, and the needle was left in place for 4 minutes before being slowly retracted. The incision was closed using silk sutures and VetBond. Mice were given 500 μL of sterile saline following surgery and returned to a cage with a heating pad for at least 1 hour to assist with recovery. Carprofen (10 mg/kg) was given for 3 days following surgery, and incisions were monitored for infection and reopening. Following surgery, we waited at least 2 weeks for viral expression before the start of behavioral experiments and drug exposure.

### Drug exposure

Morphine hydrochloride and fentanyl citrate were dissolved in sterile saline (0.9%) and delivered subcutaneously either by bolus injection or infusion using osmotic minipumps (Alzet Model 2001). Morphine solutions for minipumps were adjusted for individual body weight to achieve a dose of 63.2 mg/kg/day, and warmed to 37°C to reduce the temporal delay of drug administration. Minipumps were filled with 300 μL of solution and stored at 37°C until implantation. Before surgically implanting the pumps, mice were anesthetized with isoflurane (3-5%) and treated with carprofen (10 mg/kg, s.c.). A small section of the rump was shaved and sterilized using isopropyl alcohol and iodine before making a small incision and inserting blunt-end scissors to clear the underlying fascia. The minipump was then inserted (with the drug delivery portal oriented rostrally) into the cleared space. The incision was closed with 2-3 wound clips and swabbed with iodine for further sterility. Carprofen (10 mg/kg) was administered for 3 days following pump implantation.

Morphine exposure was interrupted using naloxone (0, 1, or 3.2 mg/kg, s.c.) dissolved in sterile saline (0.9%) twice per day and 2 hours apart, as previously described (Lichtblau and Sparber, 1981; Lefevre et al., 2020, 2023). Injections were performed between 9000h and 12000h. Behavioral assessments were conducted before drug exposure (day 0), 24 hours after pump implantation and before the first naloxone injection (day 1), or 24 hours after the last naloxone injection (day 7). The body weights of mice were tracked throughout the weeklong morphine studies and recorded before the first naloxone injection and again 15 minutes after the second injection.

### Locomotor behaviors

Locomotion was measured as previously described (Lefevre et al., 2020; Brandner et al., 2023). In brief, open-field chambers (ENV-510, Med Associates) were made from clear plexiglass and stored within sound-attenuating boxes with white noise generated by ventilation fans. The location and movement of the mouse were identified using arrays of infrared beams and recorded in real-time using Activity Monitor software (v7.0.5.10, Med Associates). Mice were habituated to the room with the open-field chambers for one hour before testing. Locomotor test sessions lasted 60 minutes for the morphine pump/injection studies, and 90 minutes for the fentanyl injection studies. Mice were placed into the center of the chamber immediately after receiving injection of morphine or lower fentanyl doses (0.063-0.2 mg/kg). For higher doses of fentanyl (0.4-0.63 mg/kg), there was a 15 minute delay between injection and placement in the open-field chamber.

### Immunohistochemistry to confirm virus expression

Mice were deeply anesthetized using sodium pentobarbital (390 mg/kg i.p, Fatal-Plus, Vortech Pharmaceuticals), and anesthetic depth was confirmed with a hard toe pinch. Perfusions were performed through the heart using ice-cold PBS (in mM: 137 sodium chloride, 2.7 potassium chloride, 11.9 phosphate buffer) until the liver cleared, followed by ∼200 mL of room temperature fixative (4% paraformaldehyde with 0.2% picric acid in 0.4M PBS, pH 6.9). Brains were extracted and post-fixed at room temperature for 1-2 hours, then placed in 10% sucrose for at least 24 hours. Brains were placed in PBS and cut once coronally posterior to the NAc using a double-edged blade. Brains were sectioned coronally into 60 μm slices using a microtome (Leica SM 2400) set to −20°C. Slices were stored in cryoprotectant composed of (in M): 5.364 ethylene glycol, 0.876 sucrose, 0.05 phosphate buffer. Every fourth section was used to identify the injection sites. When native tdTomato fluorescence was not visible, tissue was stained for tdTomato using free-floating immunohistochemistry. Sections previously stored in cryoprotectant were washed in PBS (3 x 20 minutes) and then were preabsorbed in blocking buffer (PBS containing 0.3% Triton-X 100, 1% BSA, 1% normal donkey serum, and 0.01% sodium azide) on a shaker for 3-5 hours at 4°C. Slices were incubated with agitation in primary antibody (1:2000, Living Colors® DsRed Polyclonal Antibody, Cat. #632496, Takara Bio) diluted in blocking buffer for 48 hours at 4°C. Slices were rinsed in PBS for a minimum of 4 hours and 4 washes, then incubated in secondary antibody (1:300, donkey anti-rabbit-Cy3, #711-165-152, RRID AB_2307443, JacksonImmuno) overnight at 4°C on a shaker. Slices were again washed in PBS for a minimum of 4 hours and 4 washes before being mounted on gel-coated microscope slides. Samples were protected from light and left to dry for 24 hours, then were dehydrated using increasing concentrations of ethanol (70, 95, 100, 100%) followed by xylenes twice for 25 minutes each. Slides were kept in the second xylene wash until coverslipped with DPX. Slides were left to dry in the dark for another 24 hours before imaging. Imaging was performed using the CellSens program (Evident Scientific) with an Olympus DP28 camera and Cy3 filter. Images at four AP locations (+1.7, 1.5, 1.3, 1.18) were taken based on anatomical markers (Paxinos and Franklin, 2001). Cell body fluorescence as well as any fluorescent haze in and around the NAc, including the anterior commissure, was mapped at these four locations.

### Experimental Design and Statistical Analysis

Electrophysiology experiments used hemizygous BAC transgenic mice on a C57BL/6J genetic background (Chan et al., 2012; Nelson et al., 2012) to avoid potential adverse effects of breeding to homozygosity (Kramer et al., 2011). For behavior experiments, VGF^fl/fl^ mice in a given cage were randomly assigned to receive different virus treatments (Cre or Flp) and different drug treatments (naloxone and saline). On rare occasions when it became necessary to house mice individually, data were compared to group-housed mice and combined when no difference was apparent. Analysis of variance (ANOVA) was performed using GraphPad Prism 10 and IBM SPSS Statistics v30, with factorial ANOVA models used to evaluate interactions between variables. The Greenhouse–Geisser correction was applied as needed to adjust for violations of sphericity. Significant interactions were further examined by analyzing simple effects within each level. Significant main effects were followed by Fisher’s LSD post-hoc tests. Type I error was set at α = 0.05 (two-tailed) for all comparisons. In figures, significant simple effects or post-hoc comparisons between groups are denoted by asterisks (*p < 0.05, **p < 0.01, ***p < 0.001, ****p < 0.0001). Complete statistical results, including all main effects and interactions, are provided in **Table S1**. Both female and male mice were used in all experiments, and sex was incorporated as a factor in statistical analyses. Data are presented as mean ± SEM, with individual data points from male and female mice shown as closed and open symbols, respectively.

## RESULTS

### VGF is highly expressed in NAc interneurons and medium spiny neurons

We first investigated whether VGF in the NAc is expressed in a cell type-specific manner, using published datasets from single-cell and single-nucleus RNA sequencing of tissue from NAc (Avey et al., 2018; Savell et al., 2020; Chen et al., 2021; Tran et al., 2021) as well as dorsal striatum (Saunders et al., 2018). In all datasets, VGF was enriched in neurons relative to non-neuronal populations (**Fig. 1**). Within neuronal populations, high VGF expression and prevalence were observed in inhibitory interneurons (especially somatostatin-expressing interneurons), as well as Drd1- and Drd2-expressing medium spiny neurons (MSNs). This pattern was apparent across species ranging from mice (Avey et al., 2018; Saunders et al., 2018; Chen et al., 2021) to rats (Savell et al., 2020) to humans (Tran et al., 2021), indicating conservation of VGF expression pattern in the NAc.

**Figure 1.**
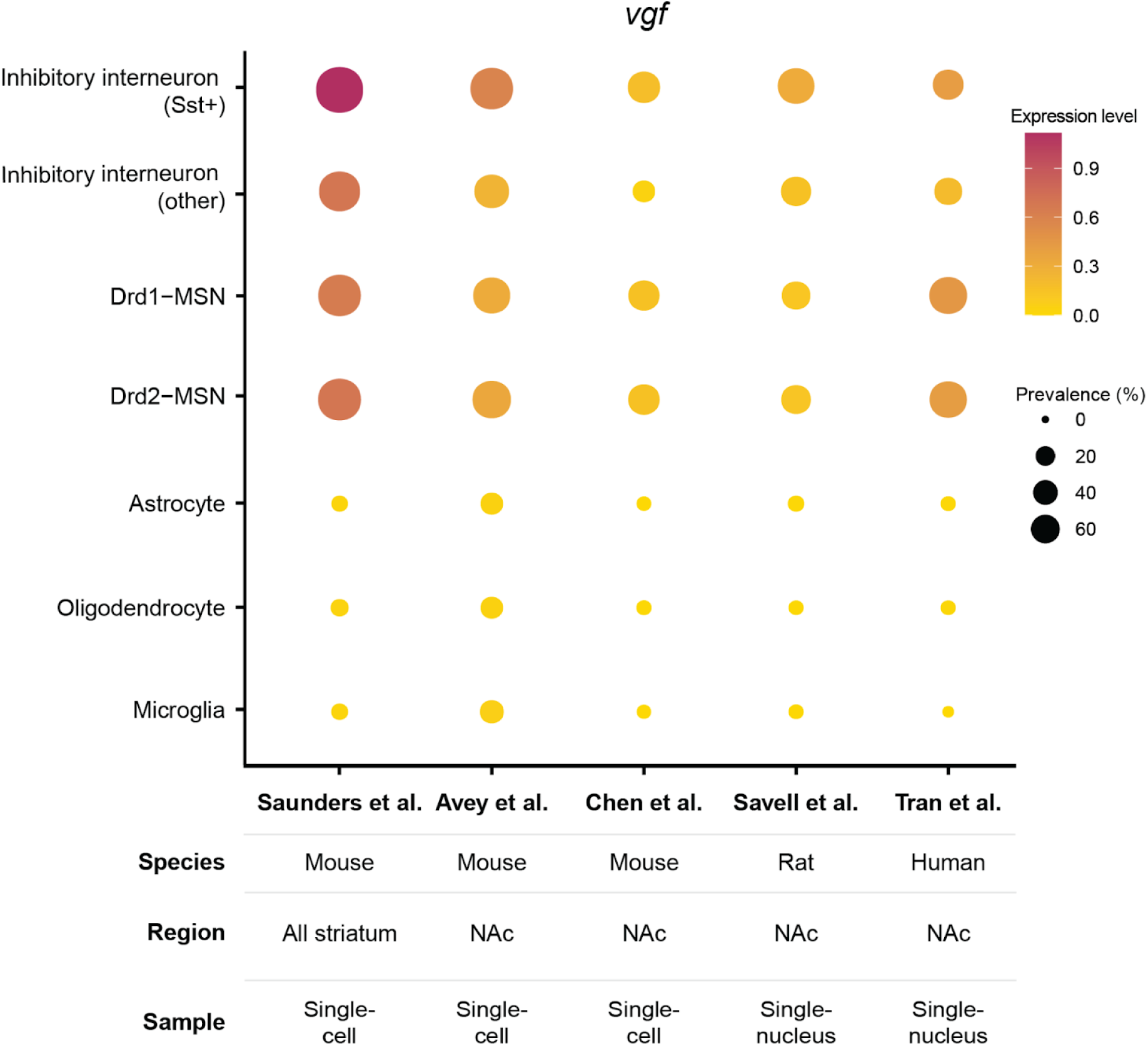
Meta-analysis of VGF expression pattern from RNA sequencing datasets. The dot plot depicts VGF expression across cell types shared by the datasets. Color intensity signifies the relative expression level of VGF, and circle size reflects the prevalence of VGF expression within each cell type. The species, region, and sample for each study are summarized below the dot plot.

### VGF expression by somatostatin interneurons across the NAc

Guided by the results of our RNA sequencing meta-analysis, we next aimed to confirm the identity and spatial location of cells with high VGF expression within the NAc. Using RNA fluorescent *in situ* hybridization, we labeled VGF and somatostatin (Sst) transcripts (**Fig. 2A**) to quantify co-localization and expression across >37,000 cells from 8 mice. Overall, we found that 47.4% of DAPI-stained cells within the NAc contained VGF transcripts (**Fig. 2B**). This proportion was similar across the NAc core, shell, or across the rostrocaudal length of the NAc (**Fig. S1**). Of all DAPI-labeled cells, 6% contained Sst transcript: 4.1% of these cells were co-labeled with VGF and Sst, while 1.9% expressed Sst alone (**Fig. 2B**). The proportion of VGF-expressing cells was higher within the Sst+ (68.2%) compared to the Sst-subset (46.1%) (**Fig. 2C**; χ² = 414.2, p < 0.0001). There was also a greater proportion of cells that are highly enriched for VGF expression in Sst+ cells: 23.2% contained a high number of VGF mRNA transcripts, compared to only 8.6% of Sst-cells (**Fig. 2C**; χ² = 520.0, p < 0.0001). From these results, we concluded that VGF has enriched expression in Sst+ cells.

**Figure 2.**
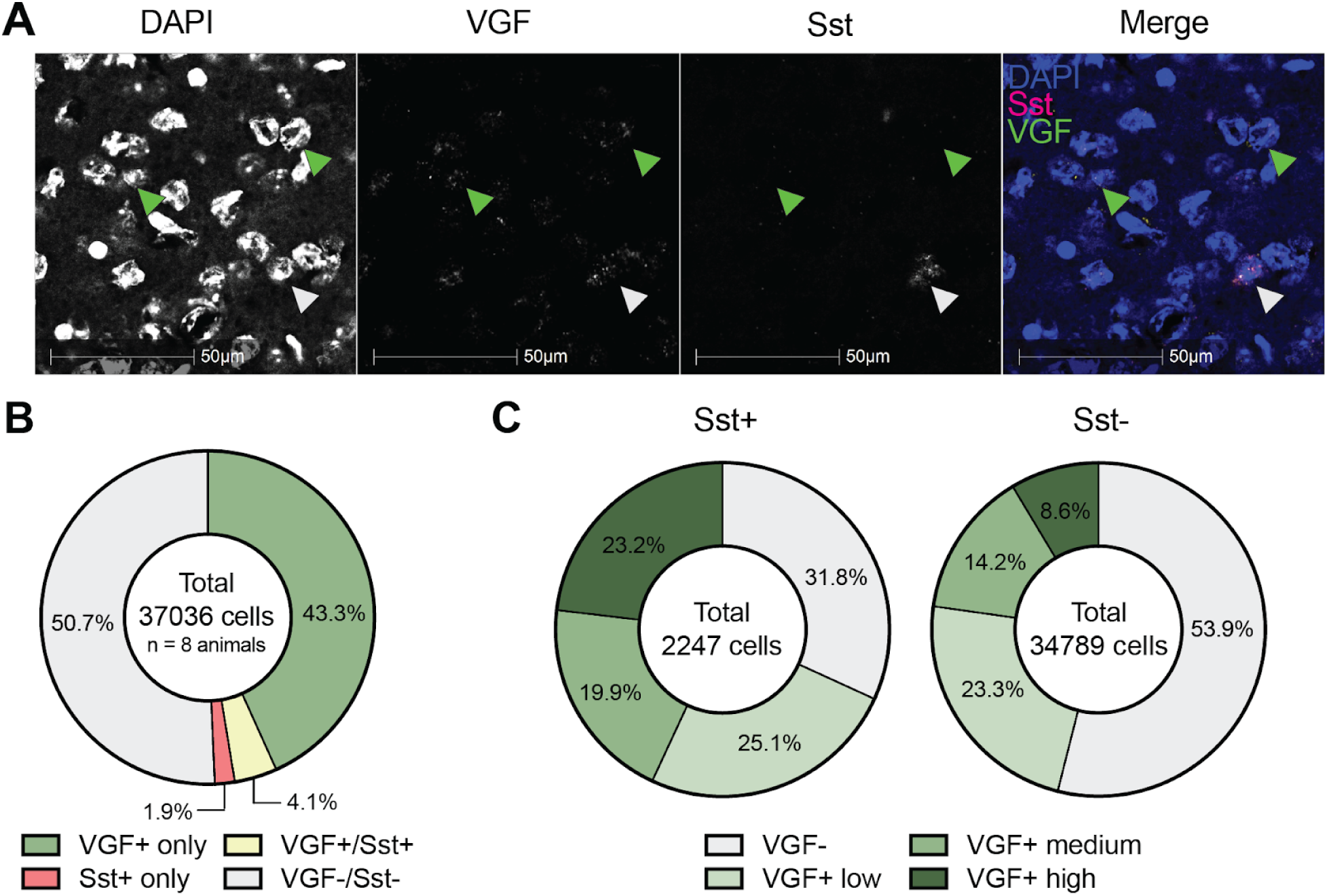
VGF co-expression by Sst cells in the nucleus accumbens. **(A)** Representative FISH images showing (from left to right) DAPI, VGF, Sst, and the merged channels. Green arrows indicate cells expressing VGF alone, and the white arrow indicates a cell co-expressing VGF and Sst. **(B)** Quantification of expression patterns across 37,036 cells imaged from 8 mice (4M/4F). **(C)** Quantification of VGF expression levels in Sst+ versus Sst-cells.

### VGF expression by Drd1- and Drd2-expressing cells

VGF expression was also high in MSNs (**Fig. 1**), which likely explains the large number of VGF+ cells that did not co-express Sst (43.3% of DAPI-labeled cells). To assess the expression and prevalence of VGF by each MSN subtype, we measured VGF co-expression with Drd1 and Drd2 transcripts (**Fig. 3A**) in a distinct sample of >27,000 cells from 6 mice. We quantified the percentage of all DAPI-labeled cells that expressed each possible combination of VGF, Drd1, and Drd2 (**Fig. 3B**). Drd1+ cells co-labeled with VGF accounted for 17.9% of all cells, compared with the 3.9% of all cells that were positive for both VGF and Drd2 (χ² = 2730, p < 0.0001). A high proportion of cells that were co-labeled with Drd1 and Drd2 were also positive for VGF (17.5%).

**Figure 3.**
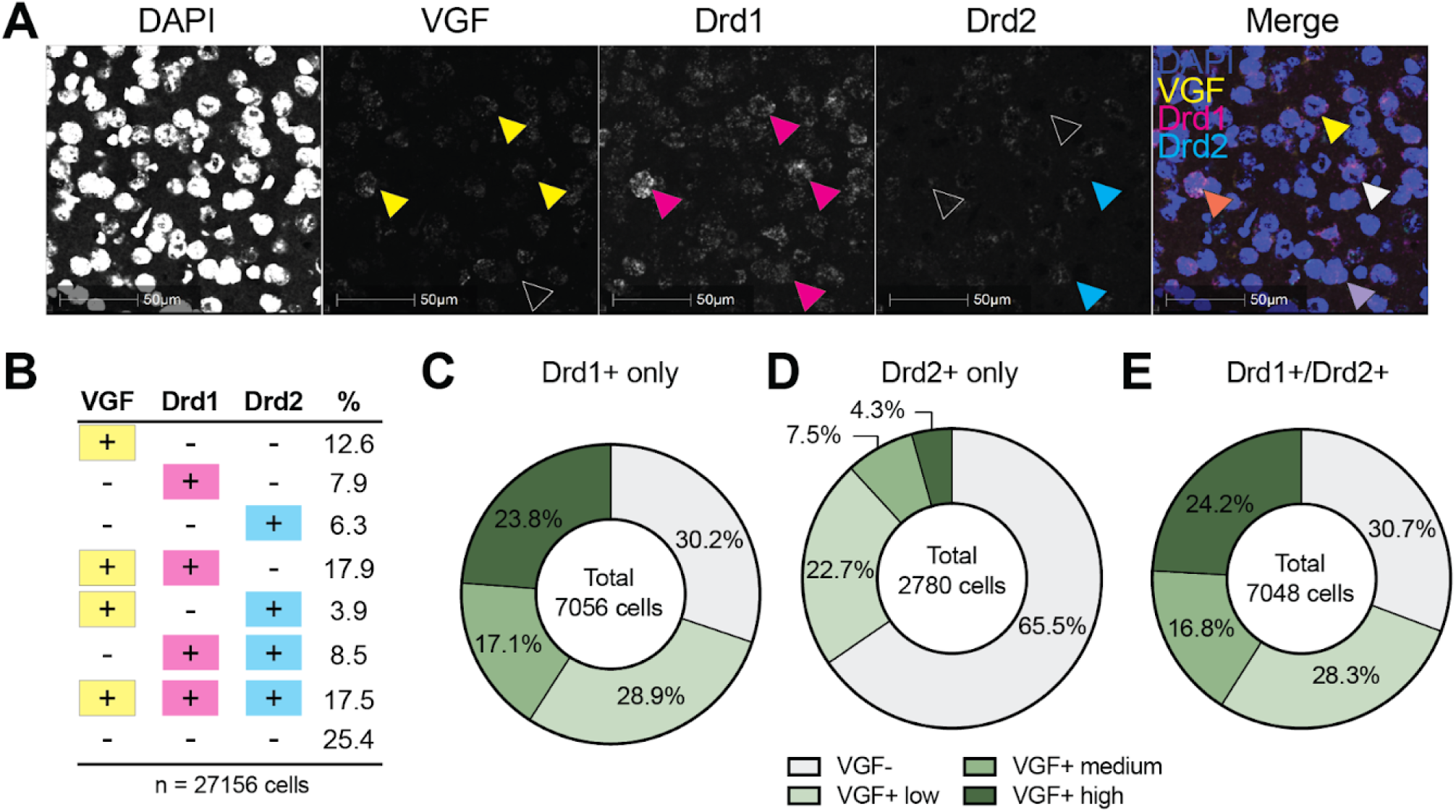
VGF co-expression in Drd1 and Drd2 cells in the nucleus accumbens. **(A)** Representative FISH images showing (from left to right) DAPI, VGF, Drd1, Drd2, and the merged channels. Yellow arrows indicate VGF+ cells, magenta arrows indicate Drd1+ cells, cyan denotes Drd2+. Black arrows indicate the absence of signal in a given channel. **(B)** Quantification of VGF, Drd1, and Drd2 expression across 27,156 cells from 6 mice (3M/3F). **(C-E)** VGF transcript levels in Drd1+ **(C)**, Drd2+ **(D),** and Drd1+/Drd2+ co-expressing cells **(E)**.

We also quantified the expression level of VGF transcripts within cells that were positive for Drd1, Drd2, or both. In cells positive only for Drd1 (**Fig. 3C**), 69.8% of Drd1+ cells also expressed VGF, compared with only 34.5% of cells positive only for Drd2 (**Fig. 3D**; χ² = 1010, p < 0.0001). More Drd1+ cells expressed high levels of VGF (23.8%) compared with Drd2+ cells (4.3%; χ² = 504.7, p < 0.0001), indicating that Drd1+ cells are enriched with VGF transcripts. The pattern of expression in cells positive for both Drd1 and Drd2 (**Fig. 3E**) was similar to cells expressing Drd1 alone, with 24.2% of Drd1/Drd2+ cells expressing high levels of VGF transcripts. Together, these data suggest that VGF is preferentially expressed in cells expressing Drd1 compared with those expressing only Drd2, and that VGF transcripts are enriched in both Drd1+ and Drd1/Drd2+ cells.

### Exogenous TLQP-62 decreases the amplitude of excitatory postsynaptic currents in MSNs

Given the abundant expression of VGF in the NAc, we next aimed to identify the effect of a VGF-derived peptide on synaptic transmission in the NAc. We focused on TLQP-62, a proteolytic product of full-length VGF with known biological activity. Using western blot analysis with an antibody recognizing the C-terminus of VGF (Lin et al., 2015; Skorput et al., 2018), we confirmed the presence of TLQP-62 in the nucleus accumbens (**Fig. S2**). Studies examining the physiological effects of TLQP-62 have found that it alters excitatory synaptic transmission (Alder et al., 2003; Bozdagi et al., 2008; Moss et al., 2008; Skorput et al., 2018). Using whole-cell voltage-clamp recordings in acute brain slices from transgenic mice expressing Drd1-tdTomato and/or Drd2-GFP reporter genes (**Fig. S3**), we measured miniature excitatory postsynaptic currents (mEPSCs) from NAc MSNs before, during, and after bath application of TLQP-62 (**Fig. 4A**) or a scrambled peptide control (**Fig. 4B**).

**Figure 4.**
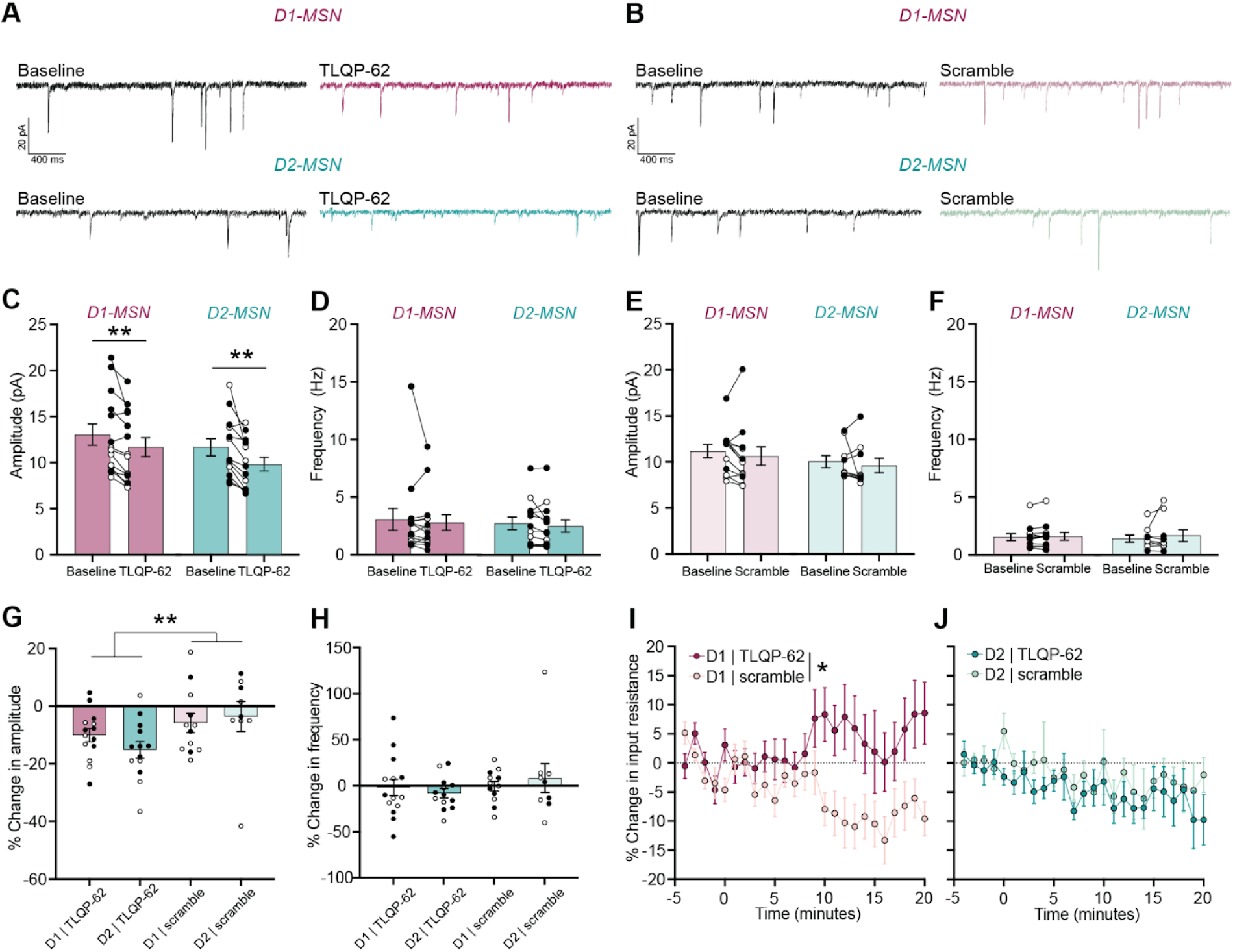
Whole-cell voltage-clamp recordings of MSNs during TLQP-62 application. **(A, B)** Example traces of mEPSCs in D1-MSNs (n = 26 cells) and D2-MSNs (n = 22 cells) before and after **(A)** TLQP-62 (50 nM) or **(B)** scrambled peptide (50 nM) bath application. **(C, D)** Average amplitude **(C)** and frequency **(D)** of mEPSCs in MSNs before and after TLQP-62 application. **p < 0.01; paired t-test. **(E, F)** Average amplitude **(E)** and frequency **(F)** of mEPSCs in MSNs before and after scrambled peptide application. **(G, H)** Average change in amplitude **(G)** and frequency **(H)** separated by cell type and peptide. **p = 0.0047; main effect of Peptide. **(I, J)** Change in the input resistance throughout the recording in D1-MSNs **(I)** and D2-MSNs **(J)**. *p = 0.0193; Time x Peptide interaction, mixed-effects model. Open and closed circles represent cells recorded from female and male mice, respectively.

Following 10 minutes of TLQP-62 application, mEPSC amplitude was significantly decreased in both D1-MSNs (t_13_ = 3.367, p = 0.0051) and D2-MSNs (**Fig. 4C**; t(12) = 3.787, p = 0.0026). However, there were no changes in mEPSC frequency in either cell type (**Fig. 4D**). The scrambled peptide control had no effect on mEPSC frequency or amplitude (**Fig. 4E-F**). TLQP-62 reliably caused a greater decrease in mEPSC amplitude than the scrambled peptide in both cell types, as evidenced by further analysis of percent change in mEPSC amplitude (**Fig. 4G**; main effect of Peptide: F_1,40_ = 8.982, p = 0.0047). There were no effects of either peptide in either cell type on percent change in mEPSC frequency (**Fig. 4H**). These results indicate that TLQP-62 decreases mEPSC amplitude for both D1- and D2-MSNs in the NAc.

We also monitored the input resistance of MSNs throughout the duration of mESPC recording, and noted that D1-MSNs showed increased input resistance 10 minutes after the TLQP-62 application, which was significantly different than scrambled peptide (**Fig. 4I**; Peptide x Time interaction: F_4.715, 104.3_ = 2.543, p = 0.0354). However, TLQP-62 did not significantly alter D2-MSN input resistance over time compared to the scrambled peptide (**Fig. 4J**). Together, these data indicate some physiological effects of TLQP-62 (i.e., decreased mEPSC amplitude) are similar in both MSN subtypes, while other effects of TLQP-62 (i.e., increased input resistance) are unique to D1-MSNs.

### Conditional VGF knockout does not alter responses to interrupted morphine exposure

We have previously shown that interrupted morphine exposure causes a reversal of the psychomotor tolerance normally produced by continuous morphine administration (Lefevre et al., 2020, 2023). This is accompanied by upregulation of VGF expression in the NAc (Lefevre et al., 2020), an effect also observed following intravenous self-administration of cocaine (Wimmer et al., 2019). We hypothesized that this upregulation of VGF in the NAc may contribute to behavioral plasticity evoked by drug exposure. We used conditional VGF knockout mice (VGF^fl/fl^) (Jiang et al., 2018) and expressed Cre recombinase via stereotaxic injection of adeno-associated virus (AAV2retro-hSyn-Cre-P2A-tdTomato) bilaterally into the NAc (**Fig. 5A**). This viral serotype is expressed locally at the site of injection (NAc), and also taken up by axon terminals and transported retrogradely to their cell body of origin (Tervo et al., 2016), thus knocking out VGF and its derived peptides from neurons projecting to and within the NAc (**Fig. 5B**). Control mice received bilateral stereotaxic injection of AAV2retro-hSyn-Flp-P2A-tdTomato into the NAc, a virus expressing a different recombinase (Flp) that does not recognize loxP sequences (**Fig. S4**). After 6 weeks of virus expression, mice were implanted with osmotic minipumps for continuous morphine delivery. Separate groups received either daily injections of naloxone to “interrupt” the effect of morphine, or daily saline injections as a control. In this study, we administered naloxone at a dose of 3.2 mg/kg, which was sufficient to reverse the psychomotor tolerance normally produced by continuous morphine administration in VGF^fl/fl^ mice (**Fig. S5**).

**Figure 5.**
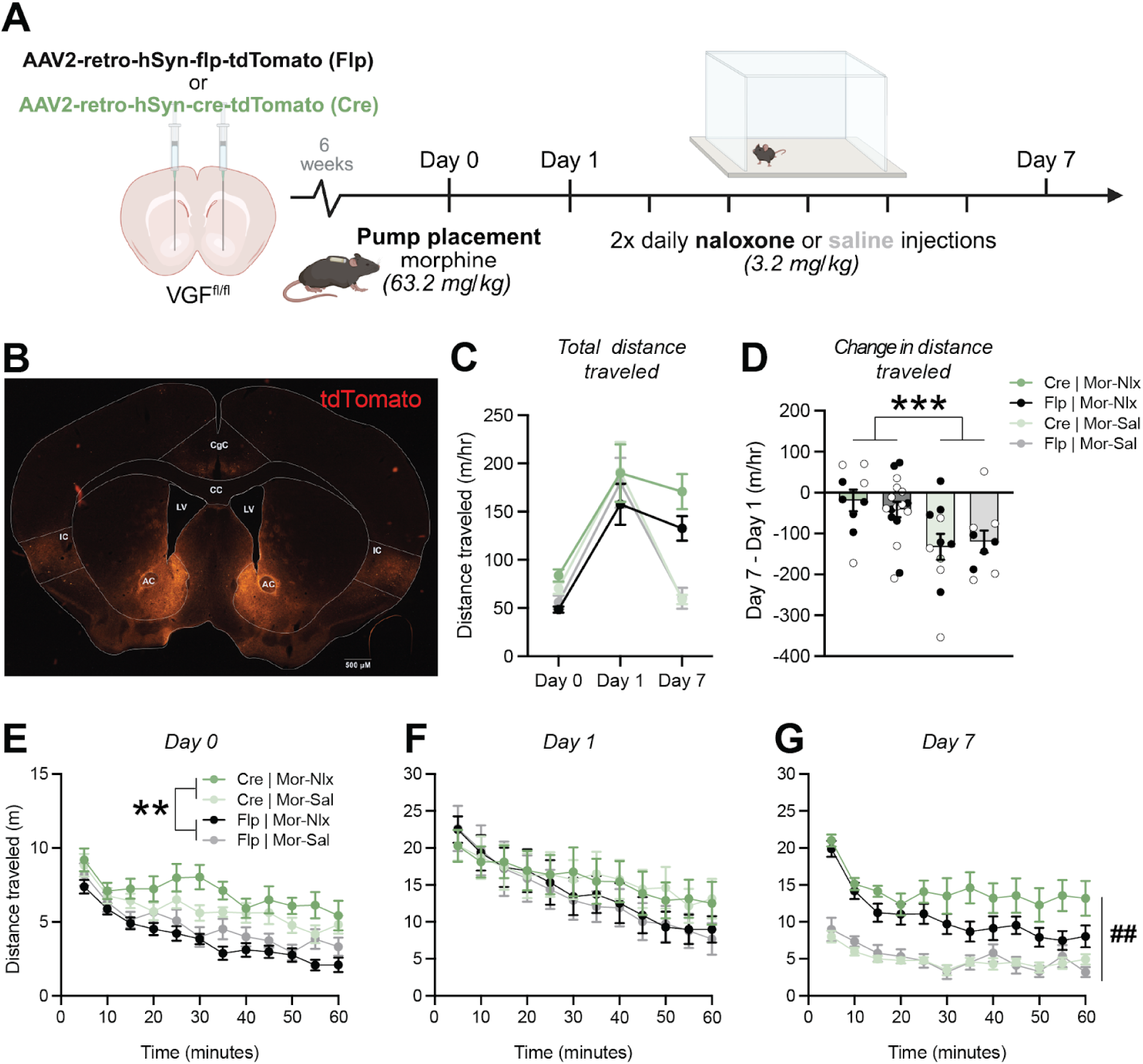
Interruption of continuous morphine exposure following conditional genetic knockout of VGF. **(A)** Experimental timeline: 6 weeks after bilateral intracranial virus injection into the NAc, mice were habituated to the open field boxes (Day 0) before osmotic pump placement. Pumps provided continuous infusion of morphine (63.2 mg/kg) interrupted twice daily (separated by 2 hours) with naloxone (3.2 mg/kg) or saline. Locomotion was measured after ∼24 hours of morphine exposure, prior to naloxone or saline administration (Day 1), and again ∼24 hours after the last naloxone injection (Day 7). **(B)** Representative image of AAV2-retro-hSyn-cre-tdTomato (Cre) injection site in the NAc and retrograde labeling of cortical regions projecting to NAc. **(C)** Total distance traveled before (Day 0) drug exposure and on the first (Day 1) and last (Day 7) days of morphine exposure. **(D)** Change in locomotor activity on Day 7 versus Day 1. ***p = 0.0007; main effect of Naloxone. **(E-G)** Time courses of average locomotor activity during sessions on Day 0 **(E)**, Day 1 **(F)**, and Day 7 **(G)**. **p < 0.0079; Time x Virus interaction. ^##^p = 0.0014; Time x Naloxone interaction. n = 9-17 animals/group. Open and closed circles represent female and male mice, respectively. AC - anterior commissure; LV - lateral ventricle; CC - corpus collosum; CgC - cingulate cortex; IC - insular cortex.

During a baseline test of open-field locomotor activity before morphine administration, we noted that mice injected with Cre virus were significantly more active than the control group injected with Flp virus (**Fig. 5C**; main effect of Virus on day 0: F_1,43_ = 22.34, p < 0.0001). This difference was particularly evident in the latter portions of the 60-minute test session (**Fig. 5E**; Virus x Time interaction: F_7.997, 343.9_ = 2.652, p = 0.0078). However, the psychomotor activation produced by morphine on Day 1 (after 24 hours of continuous exposure) was similar in all groups, regardless of virus injection (**Fig. 5C & F**).

In control groups receiving saline injections and thus exposed to morphine continuously, we observed a decrease in the psychomotor response on Day 7 compared to Day 1, reflecting the development of psychomotor tolerance (**Fig. 5C**). This change did not appear to differ as a function of virus injection, which was also evident in analysis of difference scores comparing the change in distance traveled on Day 7 versus Day 1 (**Fig. 5D**; main effect of Naloxone: F_1,39_ = 13.46, p = 0.0007). In groups receiving daily naloxone injections to interrupt the continuity of morphine exposure, the psychomotor response to morphine remained stable on Day 7 compared to Day 1, reflecting a reversal of psychomotor tolerance (**Fig. 5G**). However, the magnitude of this effect did not differ as a function of virus injection (**Fig. 5D**; Virus x Naloxone interaction: F_1,39_ = 1.045, p = 0.3129). We therefore concluded that conditional knockout of VGF from the NAc and its inputs does not alter locomotor behaviors elicited by continuous or interrupted morphine exposure, but surprisingly causes hyperlocomotion in a drug-naïve state.

### Conditional VGF knockout amplifies psychomotor sensitization following repeated fentanyl exposure

In a separate cohort of mice, we examined how conditional knockout of VGF from the NAc and its inputs affects sensitivity to morphine (a prototypical opioid) and fentanyl, an opioid with pressing public health relevance. This experiment was conducted using a more conventional protocol involving non-contingent drug injections. We measured sensitivity to low doses of both morphine and fentanyl after 2-6 weeks of virus expression, and then examined the development and persistence of psychomotor sensitization following repeated exposure to a higher dose of fentanyl. After injecting VGF^fl/fl^ mice with the same Cre- and Flp-expressing viruses described above, we performed long-term, cyclic injections of low-dose opioids (**Fig. 6A, S6**). Every two weeks after stereotaxic surgery, we monitored locomotor activity following injection with saline, morphine (0.2 mg/kg, s.c.), and fentanyl (0.063 mg/kg, s.c.) on consecutive days. We again observed a trend towards VGF^fl/fl^ mice injected with Cre virus becoming hyperactive at baseline in the absence of opioid exposure, but did not observe any significant differences in sensitivity to a low dose of either morphine or fentanyl at any time point (**Fig. S7**).

**Figure 6.**
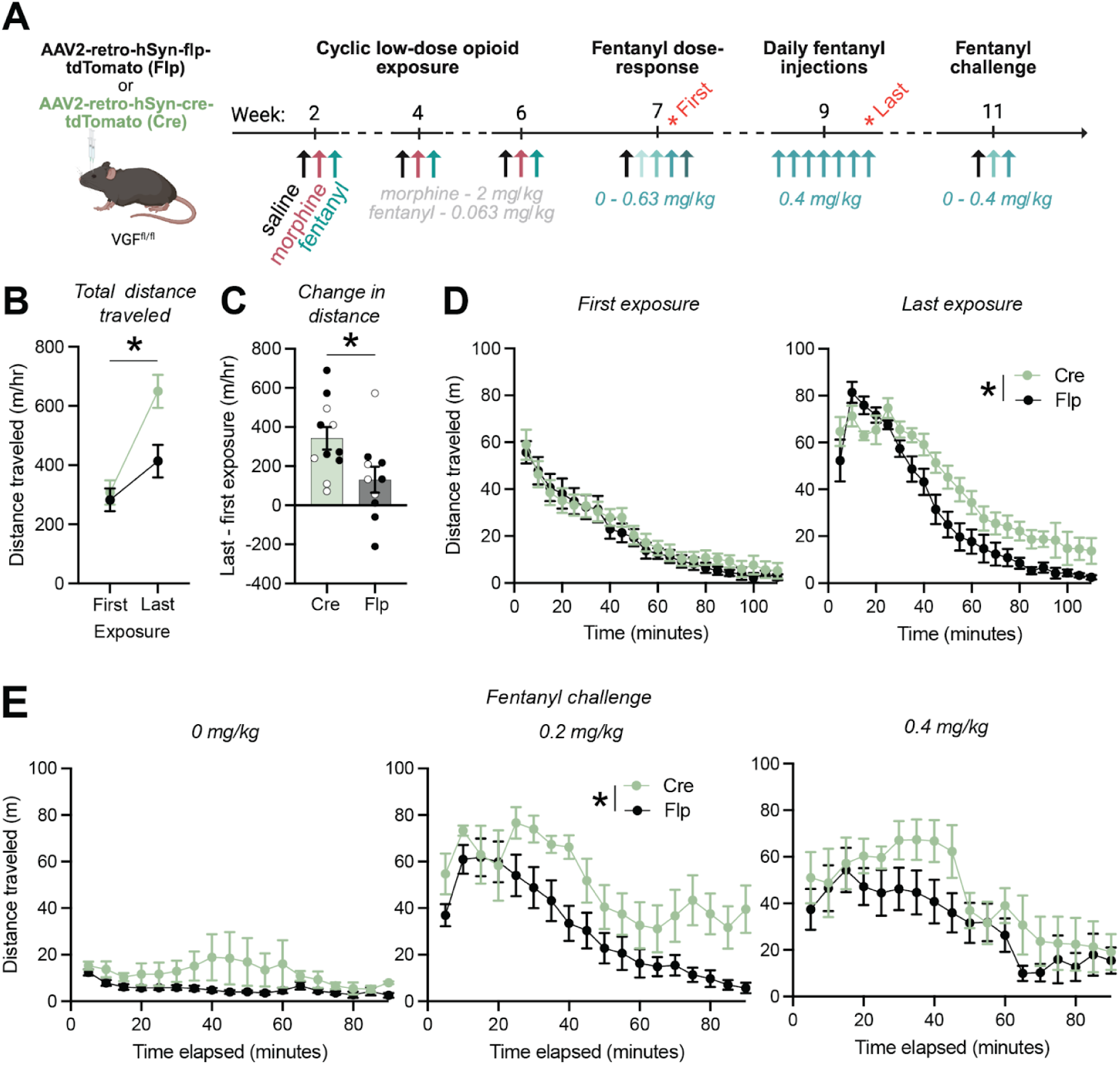
Chronic fentanyl exposure following conditional genetic knockout of VGF. **(A)** Experimental timeline: 2 weeks after bilateral intracranial virus injection into the NAc, mice were exposed to low doses of opioids. On separate days, mice received injections of saline, morphine (2 mg/kg), and fentanyl (0.063 mg/kg), with locomotion recorded after each injection. The same sequence was repeated 4 and 6 weeks after viral delivery. At week 7, mice underwent a fentanyl dose-response series across five days, followed at week 9 by daily high-dose fentanyl injections (0.4 mg/kg) for 7 days. Two weeks after the final injection, mice received a fentanyl challenge consisting of increasing doses on separate days. **(B)** Total distance traveled on the first exposure to 0.4 mg/kg fentanyl during the week 7 dose–response session and the last exposure during week 9. *p = 0.043, Day x Virus interaction. **(C)** Change in locomotor activity between the first and last 0.4 mg/kg fentanyl exposures. *p = 0.043, main effect of Virus. **(D)** Locomotor time course during from the first (left) and last (right) exposure to 0.4 mg/kg fentanyl. *p = 0.0007, Time x Virus interaction. **(E)** Fentanyl challenge was conducted across three days with doses of 0 (left), 0.2 (middle), and 0.4 mg/kg (right), shown as within-session time courses. *p = 0.0233, main effect of Virus. n= 10-11 animals/group. Open and closed circles represent female and male mice, respectively.

Seven weeks after virus injection, we performed an acute dose-response study using escalating doses of fentanyl injections (0-0.63 mg/kg, s.c.) on consecutive days (**Fig. 6A**). However, we did not observe any significant differences between groups in acute fentanyl sensitivity (**Fig. S8B & C**). Nine weeks after virus injection, we exposed the same group of mice to daily injection of a high dose of fentanyl (0.4 mg/kg, s.c.) for 7 day to generate psychomotor sensitization. While both groups showed similar acute sensitivity in terms of their first exposure to 0.4 mg/kg fentanyl, the development of psychomotor sensitization was amplified in the Cre group compared to the Flp group (**Fig. 6B**; Virus x Day interaction: F_1,17_ = 4.97, p = 0.04). This phenotype was also apparent after computing the change in distance travelled between Days 1 and 7 for each individual animal (**Fig. 6C**; F_1,17_ = 4.779, p = 0.0431), and most evident during the latter portions of the Day 7 test session (**Fig. 6D**; Virus x Time interaction: F_21,315_ = 2.3856, p = 0.0007). The amplification of sensitization persisted after two weeks of abstinence, as evidenced by a heightened response to challenge with a moderate dose of fentanyl (0.2 mg/kg) (**Fig. 6E**). Given these data, we concluded that VGF-derived peptides acting in the NAc may dampen psychomotor sensitization following daily fentanyl exposure, providing a brake that blunts drug-evoked plasticity.

## DISCUSSION

Dissecting the mechanisms by which neuropeptides influence dysregulated synaptic plasticity and NAc-related behaviors could identify new, precise targets for addiction treatment. Prior work has established VGF-derived peptides as regulators of plasticity in the spinal cord (Skorput et al., 2018), neocortex (Selten et al., 2025), and hippocampus (Thakker-Varia et al., 2007; Bozdagi et al., 2008; Lin et al., 2015), in both adaptive (recovery after nerve injury, memory formation) and maladaptive states (persistent hypersensitivity/chronic pain, depression). Past work has shown that, in the NAc, VGF is upregulated following cocaine self-administration (Wimmer et al., 2019) and specific patterns of morphine exposure (Lefevre et al., 2020), suggesting that it may play a role in drug-evoked plasticity. VGF and its derived peptides therefore pose an intriguing potential regulator of reward neurocircuitry. Here, we characterized VGF expression in the NAc, examined its contribution to synaptic transmission onto D1- and D2-MSNs, and assessed its function in models of opioid exposure and withdrawal.

NAc cell types are diverse, and their corresponding functions and influence on downstream targets differ widely depending on their transcriptional signature and cell markers (Lobo and Nestler, 2011; Ribeiro et al., 2018; Tran et al., 2021; Gallegos et al., 2023). Prior studies using single-cell or nuclei RNA-sequencing showed that, across species, VGF is widely expressed by NAc neurons, especially inhibitory interneuron and MSN populations. Our FISH studies corroborated and expanded on these findings. VGF expression was similar across the NAc core and shell subregions, as well as across the rostrocaudal length. Both Sst+ interneurons and D1-MSNs had high VGF expression, while D2-MSNs had a lower level of VGF expression. In cells that expressed both Drd1 and Drd2, the level of VGF expression was high and comparable to cells that express only Drd1. These data suggest that across MSN subtypes within the NAc, VGF expression is tightly linked with Drd1 expression.

VGF-derived peptides, particularly the C-terminal peptide TLQP-62, are known to facilitate plasticity within the central nervous system, but the exact mechanism by which they act is unknown. Prior studies examining the effect of TLQP-62 on synaptic transmission and physiology have employed calcium imaging and electrophysiology in acute brain or spinal cord slices or cultured neurons (Alder et al., 2003; Bozdagi et al., 2008; Moss et al., 2008; Skorput et al., 2018). These papers showed a range of effects of TLQP-62, differing based on methodology and preparation, and generally found alterations in excitatory activity following TLQP-62 exposure. We observed a decrease in mEPSC amplitude across both D1- and D2-MSNs, which was reliably greater with active versus control peptide. This effect, coupled with the lack of change in mEPSC frequency, points to a postsynaptic change in glutamatergic signaling caused by TLQP-62.

We previously reported that interrupted morphine administration produced both psychomotor sensitization and upregulated expression of VGF in the NAc, with neither effect observed following continuous morphine exposure (Lefevre et al., 2020). We therefore hypothesized that VGF expression in the NAc may contribute to behavioral plasticity produced by chronic opioid administration. However, conditional knockout of VGF from the NAc and its inputs did not affect locomotor responses to either continuous or interrupted morphine. We note that in VGF^fl/fl^ mice, naloxone administration reversed the psychomotor tolerance normally produced by continuous morphine delivery, but did not induce classical psychomotor sensitization (i.e., a significant increase in the day 7 response compared to day 1). This model therefore may not have provoked the plasticity in which we had hypothesized VGF to act, thus diminishing the effects of VGF knockout. To evaluate the contribution of VGF to bona fide sensitization, we used a second model of daily fentanyl injections that robustly elicited locomotor sensitization. Conditional knockout of VGF from the NAc and its inputs resulted in marked amplification of this sensitization, suggesting that VGF and its derived peptides may normally counteract the development of psychomotor sensitization. Without full functionality of VGF, the mechanisms underlying opioid-evoked plasticity may proceed unchecked and allow for heightened psychomotor sensitization.

The intersection between our behavioral and electrophysiological findings provides ground for speculation regarding potential mechanism(s) of action for VGF and TLQP-62. Conventional models suggest that the enhancement of psychomotor sensitization we observe following conditional VGF knockout could be caused by enhanced D1-MSN activity, and/or diminished D2-MSN activity (Hikida et al., 2010; Lobo et al., 2010; Ferguson et al., 2011; Severino et al., 2020). We found that exogenous TLQP-62 decreased mEPSC amplitude in both D1- and D2-MSNs, so conditional VGF knockout might increase excitatory input to both cell types – a symmetric effect that does not easily map onto these conventional models. This contrasts with the asymmetric effect of TLQP-62 on input resistance, which increased in D1-MSNs but not D2-MSNs. Input resistance and intrinsic excitability are often positively correlated, but the implication that conditional VGF knockout would decrease D1-MSN input resistance and excitability is also inconsistent with conventional models of sensitization. This inconsistency could be reconciled if TLQP-62 increases input resistance by closing a channel with an excitatory influence on D1-MSN activity, such as a hyperpolarization-activated cyclic nucleotide-gated (HCN) channel (Maria-Rios et al., 2023). TLQP-62 is hypothesized to act via the TrkB/BDNF pathway (Lin et al., 2014, 2015; Jiang et al., 2019b, 2019a), but there is no direct evidence that TLQP-62 binds to either target, and other functionally relevant binding partners have not been identified. Therefore, ion channels could serve as TLQP-62 binding partners or downstream mediators of its effects. These possibilities could be further investigated using whole-cell current-clamp recordings from D1-MSNs, to characterize how TLQP-62 alters intrinsic excitability. We also note that hyperlocomotion in the drug-naive state following conditional VGF knockout from the NAc and its inputs may also be consistent with increased D1-MSN activity (Kravitz et al., 2010; Rothwell et al., 2014).

In contrast to our finding of basal hyperlocomotion, a prior study manipulating VGF in the NAc showed no change in locomotor behaviors after conditional knockout (Jiang et al., 2018). These disparate observations may be due to methodological differences: while Jiang et al. (2018) used a locally-targeted virus (AAV2-Cre-GFP), our studies used retrograde viruses that disrupted not just locally produced VGF, but any VGF-derived peptides that may be released from projection neurons into the NAc. Thus, it is possible that the hyperlocomotive phenotype is driven by projections into the NAc that release TLQP-62 or other VGF-derived peptides. Jiang et al. (2018) also examined the role of VGF in the NAc in chronic social defeat stress and the resulting depressive-like behaviors. Knocking out VGF in the NAc reduced social avoidance and immobility time in a forced swim test following chronic social defeat stress. Disruption of VGF in the NAc therefore counteracted the detrimental effects of chronic social defeat stress (Jiang et al., 2018). Our data suggest these effects of VGF could involve regulation of D1-MSN activity, which can influence susceptibility and resilience to chronic social defeat stress (Francis et al., 2015).

Our data provide a basis for VGF-derived peptides as a modulator of reward neurocircuitry and psychomotor sensitization. Characterizing VGF expression across the NAc revealed its presence in key cell types within the region, including D1- and D2-MSNs. Furthermore, the VGF-derived peptide TLQP-62 robustly decreased the amplitude of excitatory postsynaptic currents onto MSNs in the NAc. Finally, knocking out VGF in the NAc amplified psychomotor sensitization following chronic fentanyl exposure. While this is the first study manipulating VGF and its derived peptides within the context of opioid exposure, our findings, combined with the prior studies showing its involvement in plasticity, indicate a promising target for manipulating the detrimental effects of opioid exposure. Advancements in drug delivery could eventually allow for cell-type specific targeting via nanoparticles (Zhang et al., 2016; Sierri et al., 2024; Liu et al., 2025), and studies are already underway to assess the possibility of using VGF-derived peptides as a therapeutic target for treating diseases of the central nervous system (Arora and Singh, 2021). This raises the intriguing possibility of harnessing VGF-derived peptides for targeted therapies to treat substance use disorders and other conditions involving the dysregulation of NAc circuitry.

## ACKNOWLEDGEMENTS

We thank Galina Kalyuzhnaya and Dr. Yasushi Nakagawa for providing technical assistance and support. The viral vectors used in this study were generated by the University of Minnesota Viral Innovation Core, part of the Center for Neural Circuits in Addiction (P30 DA048742). Confocal imaging was performed with the resources and assistance from staff at the University of MInnesota University Imaging Centers (RRID SCR_020997). This work was supported by the University of Minnesota’s MnDRIVE (Minnesota’s Discovery, Research, and Innovation Economy) initiative (PER); grants from the National Institutes of Health (F30 DA060027 to APA, T32 GM008244 to APA, T32 DA007234 to APA, F31 MH133285 to RMD, and R01 DA048946 to PER); the Dr. Warren and Henrietta Warwick Fellowship (APA); and Department of Defense grant W81XWH2010509 (LV).

---

**Figure S1.**
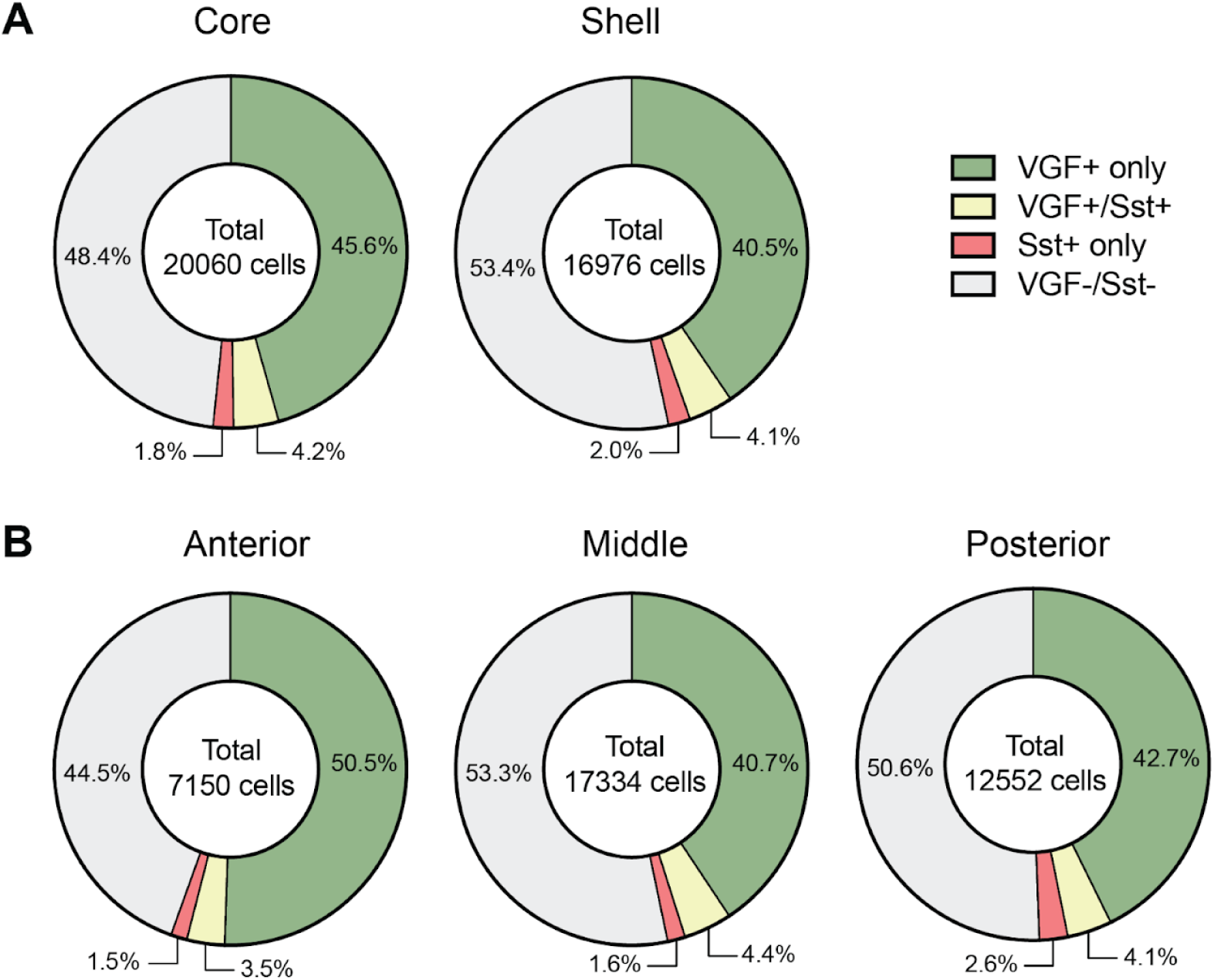
VGF and somatostatin expression across NAc subregions. **(A)** Distribution of VGF and Sst+ cells across the NAc core (left) and shell (right). (**B)** Distribution of VGF and Sst+ cells across the anterior (AP +1.54-1.98), middle (AP +1.1-1.42), and posterior (AP +0.62-0.98) regions of the NAc.

**Figure S2.**
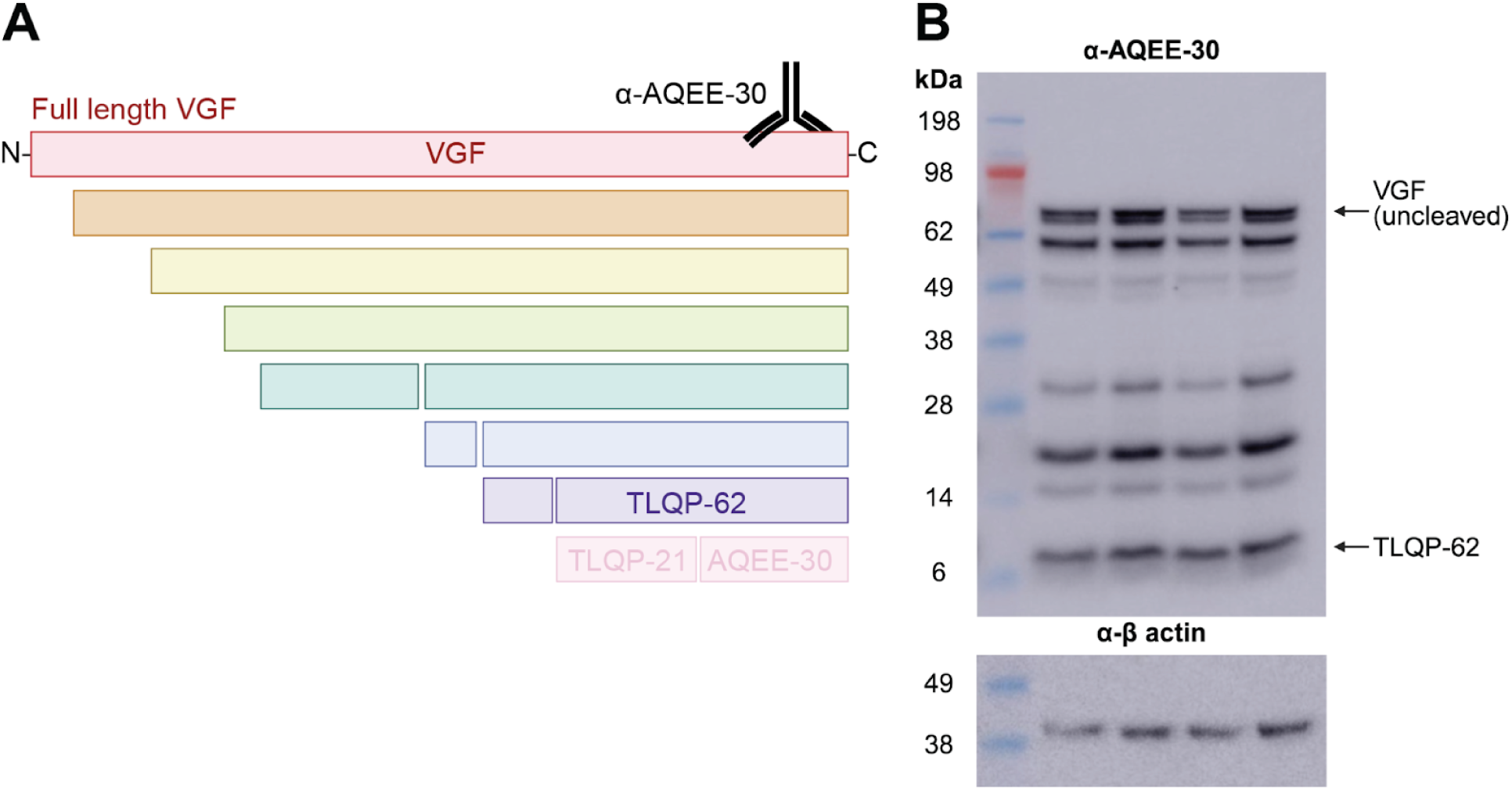
Detection of VGF-derived peptides within the NAc. **(A)** Schematic depicting the full-length VGF protein (red) and derived peptides. TLQP-62 (purple) is a ∼6 kDa C-terminal peptide. The antibody used for western blot analysis (ɑ-AQEE-30) recognizes the C-terminus of VGF. Not all VGF-derived peptides are depicted. **(B)** Western blot of samples from the NAc of 4 C57BL/6J mice (2M/2F). TLQP-62, identified by its known molecular weight, is one of several C-terminal fragments produced from the full-length VGF. β-actin served as the loading control.

**Figure S3.**
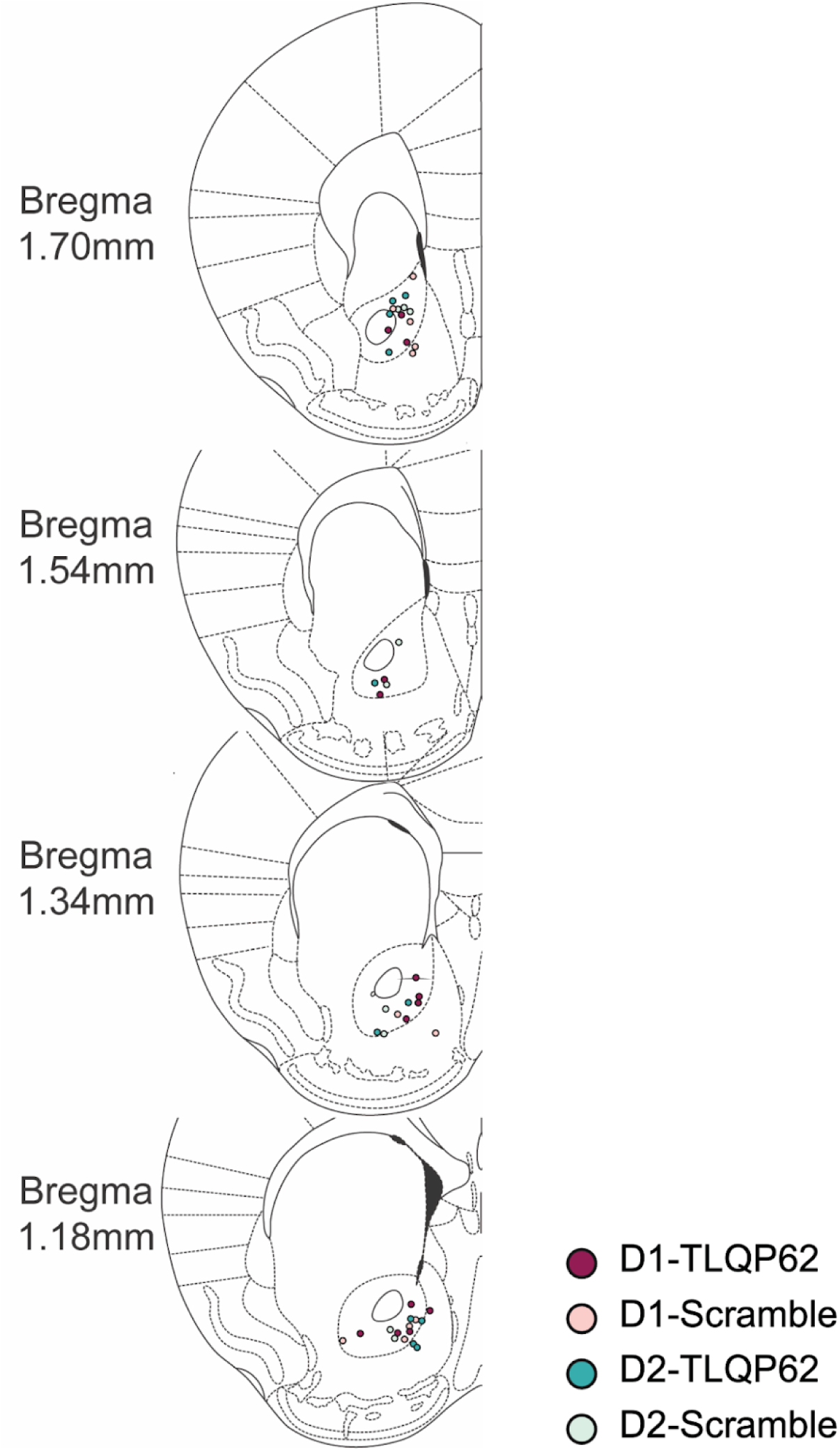
Anatomical location of whole-cell patch-clamp recordings in acute NAc brain slices. Colors define the MSN subtype and peptide treatment for each cell.

**Figure S4.**
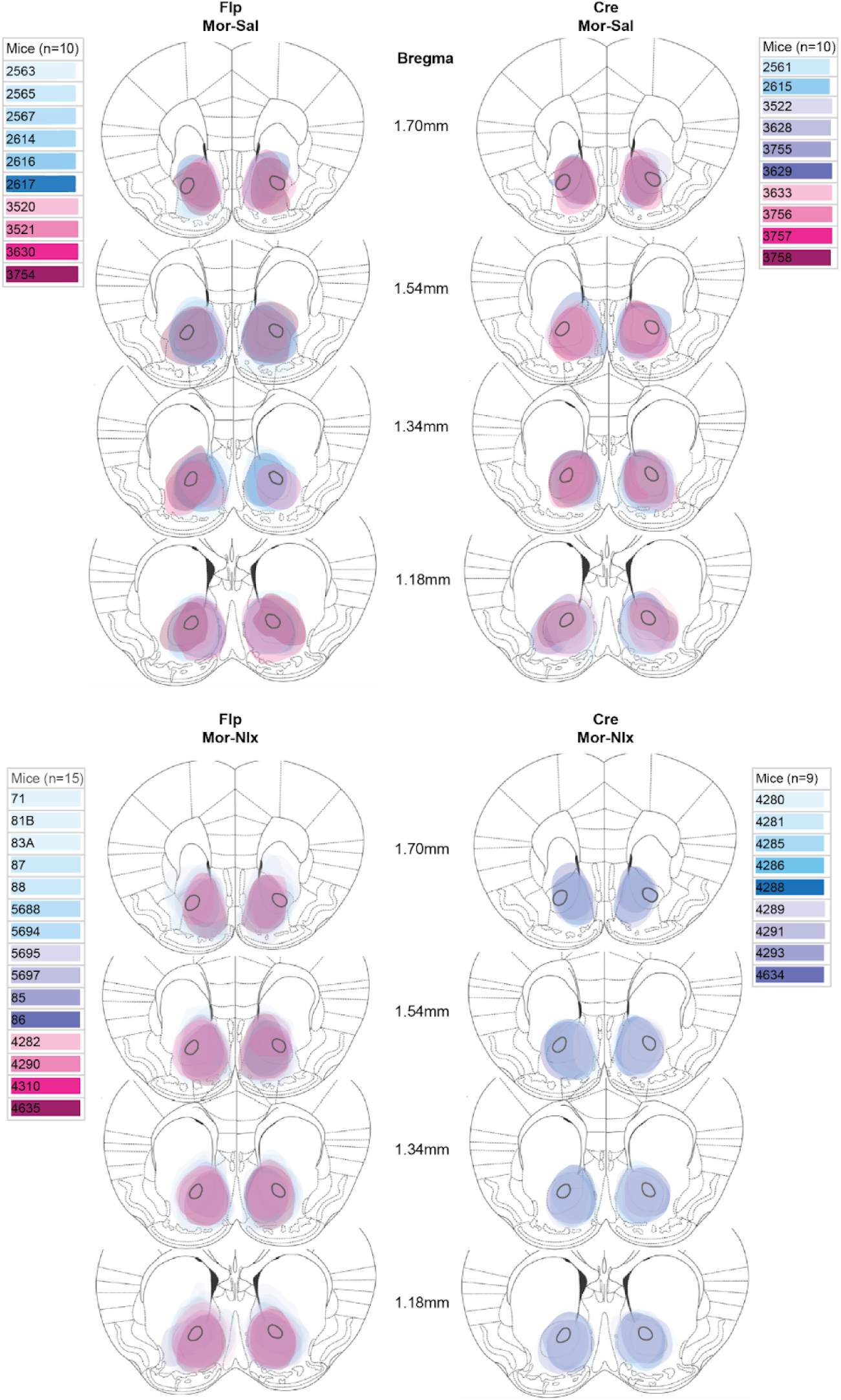
Anatomical location of virus injection sites for VGF^fl/fl^ mice exposed to interrupted morphine. Three mice are not depicted due to poor tissue quality.

**Figure S5.**
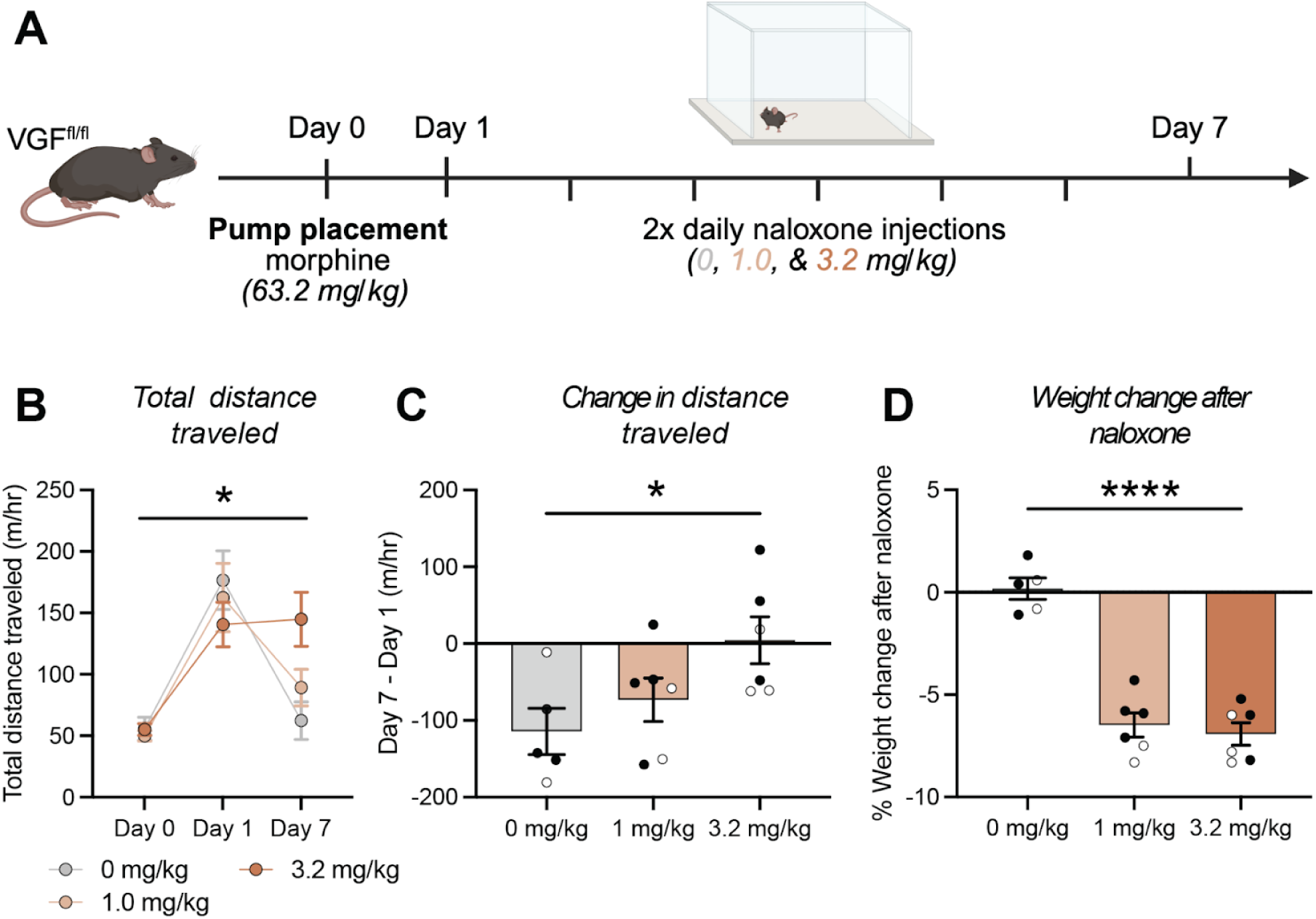
Naloxone dose-response analysis for interruption of continuous morphine exposure in VGF^fl/fl^ mice. **(A)** Experimental timeline. Mice were habituated to the open field boxes (Day 0) before osmotic pump placement. Pumps provided continuous infusion of morphine (63.2 mg/kg) interrupted twice daily (separated by 2 hours) with naloxone (0-3.2 mg/kg) or saline. Locomotion was measured after ∼24 hours of morphine exposure, before naloxone or saline administration (Day 1) and again ∼24 hours after the last naloxone injection (Day 7). **(B)** Total distance traveled before (Day 0) drug exposure and on the first (Day 1) and last (Day 7) days of morphine exposure. *p = 0.047, Day x Dose interaction. **(C)** Change in locomotor activity on Day 7 versus Day 1, as a function of naloxone dose. *p = 0.024, linear effect of Dose. **(D)** Average weight change after naloxone injections, separated by dose. *p <0.0001, main effect of Dose. (n = 5-6 animals/group). Open and closed circles represent female and male mice, respectively.

**Figure S6.**
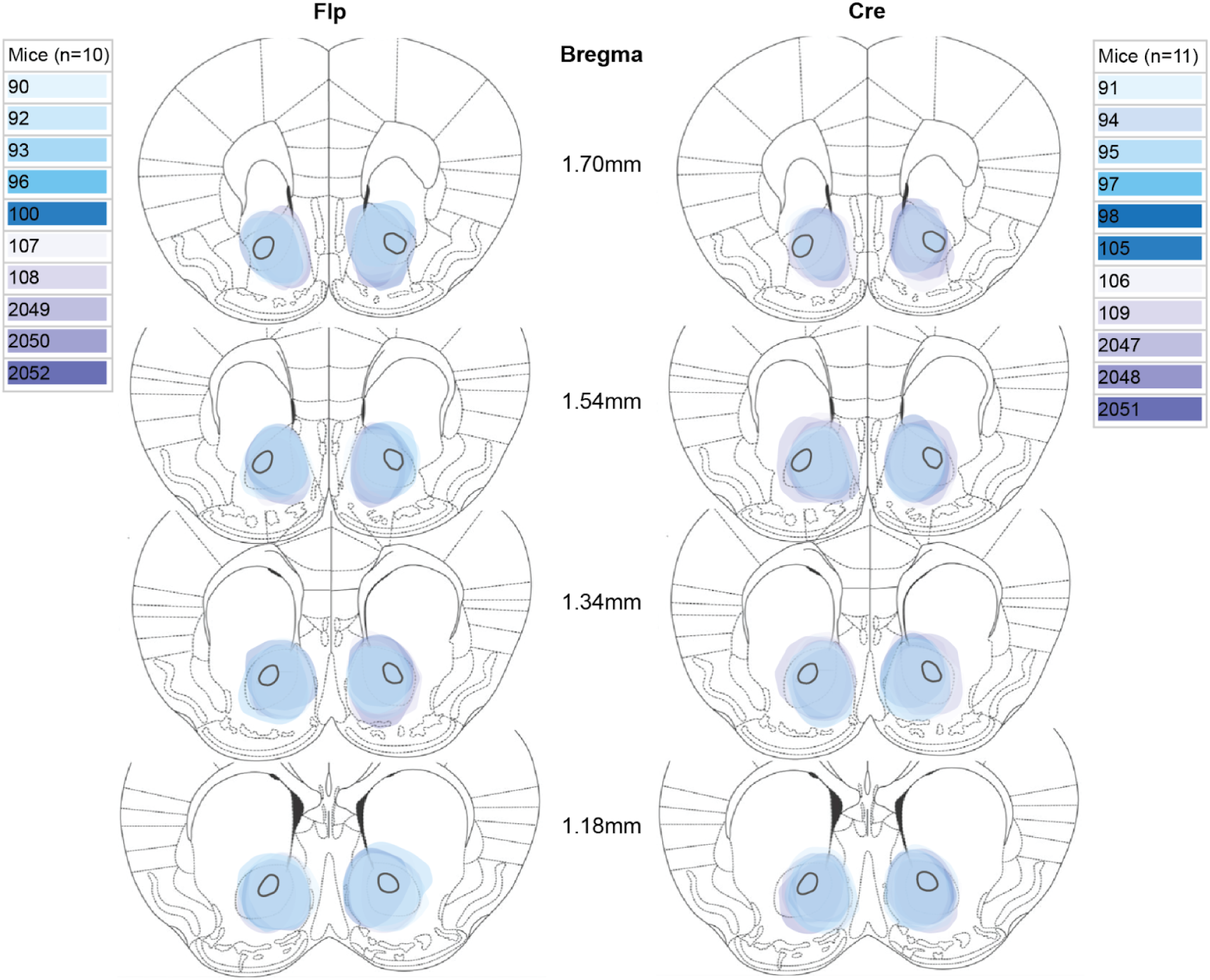
Anatomical location of virus injection sites for VGF^fl/fl^ mice exposed to chronic fentanyl.

**Figure S7.**
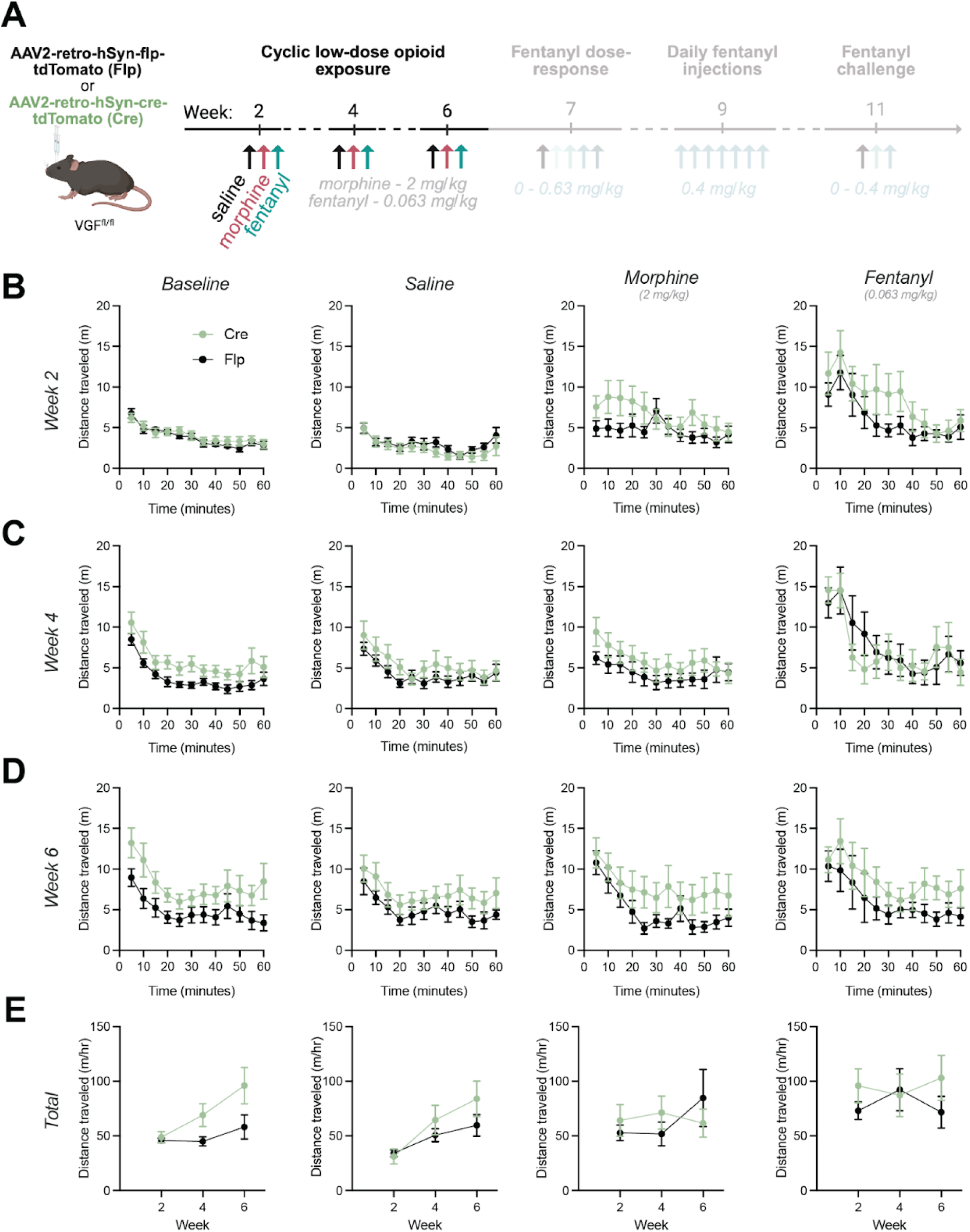
Locomotor responses to repeated low-dose opioid exposure. **(A)** Experimental timeline: 2, 4, and 6 weeks after intracranial viral injections in the NAc, sensitivity to low-dose exposure was evaluated in a cyclic fashion. Following a baseline/habituation session, mice were sequentially exposed to saline, morphine (2 mg/kg), and fentanyl (0.063 mg/kg) on separate days. **(B-D)** Locomotor activity within a single session during baseline, saline, morphine, and low-dose fentanyl at weeks 2 **(B)**, 4 **(C)**, and 6 **(D)**. **(E)** Total distance traveled across weeks 2, 4, and 6.

**Figure S8.**
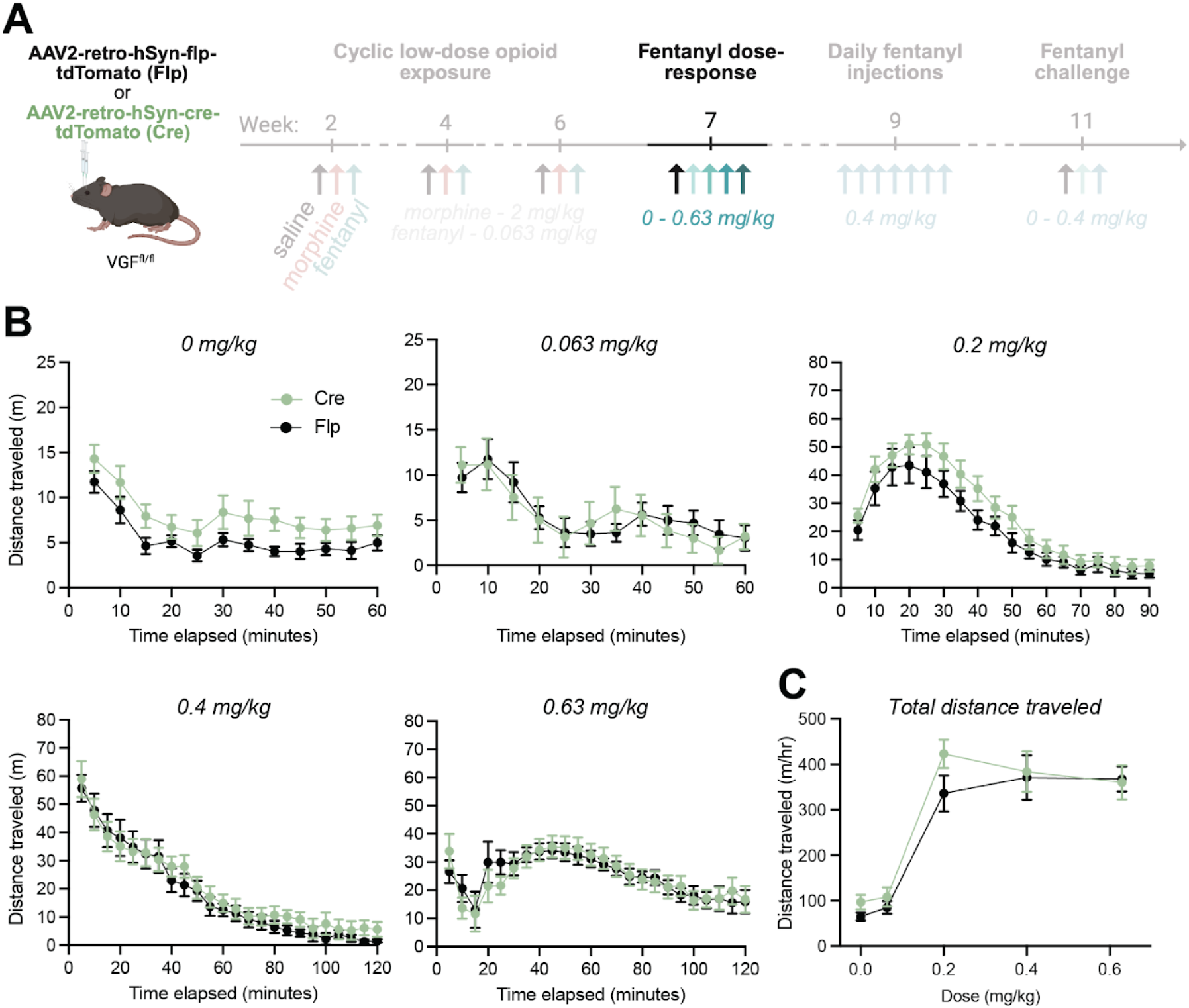
Dose–response analysis of acute fentanyl sensitivity following cyclic low-dose opioid exposure. **(A)** Experimental timeline highlighting the fentanyl dose-response during week 7. Mice received escalating fentanyl doses (0, 0.063, 0.2, 0.4, and 0.63 mg/kg) on consecutive days. **(B)** Locomotor activity within a single session at each fentanyl dose. **(C)** Distance per hour traveled at each dose.

